# Real-time single-molecule 3D tracking in E. coli based on cross-entropy minimization

**DOI:** 10.1101/2022.08.25.505330

**Authors:** Elias Amselem, Bo Broadwater, Tora Hävermark, Magnus Johansson, Johan Elf

**Author notes:** These authors contributed equally to this work (Elias Amselem, Bo Broadwater).

## Abstract

Sub-ms 3D tracking of individual molecules in living cells is an important goal for microscopy since it will enable measurements at the scale of diffusion limited macromolecular interactions. Here, we present a 3D tracking principle based on the true excitation point spread function and cross-entropy minimization for position localization of moving fluorescent reporters that approaches the relevant regime. When tested on beads moved on a stage, we reached 67nm lateral and 109nm axial precision with a time resolution of 0.84 ms at a photon count rate of 60kHz, coming close to the theoretical and simulated predictions. A critical step in the implementation was a new method for microsecond 3D PSF positioning that combines 3D holographic beam shaping and electro-optical deflection. For the analysis of tracking data, a new point estimator for diffusion was derived and evaluated by a detailed simulation of the 3D tracking principle applied to a fictive reaction-diffusion process in an *E. coli*-like geometry. Finally, we successfully applied these methods to track the Trigger Factor protein in living bacterial cells. Overall our results show that it is possible to reach sub-millisecond live-cell single-molecule tracking, but that it is still hard to resolve state transitions based on diffusivity at this time scale.

## Introduction

Advances in microscopy and nanoscopy are approaching the temporal and spatial scale where intracellular biochemistry occurs. Single-molecule tracking is a critical technique to study intracellular kinetics by analyzing changes in diffusivity[1] without perturbing the cells [2]. Recently, improved single-molecule localization for reporters with a limited photon budget was demonstrated, addressing a key limitation in live-cell single-molecule tracking[3–16]. Enhanced localization precision is here accomplished by excitation with structured optical beams and comparing the modulations in emitted fluorescence to the excitation patterns. The principle, referred to as modulation enhanced localization (SM-MEL), is different from what is traditionally done in STED [17, 18]and PALM/STORM[19, 20] where the emitted point spread function (PSF) is used for localization. Under the umbrella of SM-MEL, several strategies have recently been demonstrated, each optimized for a different purpose. The camera-based versions like ROSE[3], SIMPLE[4], SIMFLUX[5], and ModLoc[6] target a large field of view. The imaging-oriented scanning counterparts, MINFLUX[7, 8] and p-MINFLUX[9], aim for an optimal resolution with a minimal photon budget per fluorophore localization assuming slow-moving/stationary reporters. Tracking-oriented systems like Orbital scanning [10–12, 21], SMCT–FCS [13], 3D-DyPLoT [14, 15], and TSUNAMI [16] target spectroscopic and dynamical processes. Although these strategies have very different implementations, they all localize reporters using the SM-MEL principle under various assumptions. Both the camera and the minimal photon flux strategies have shown great performance in imaging; however, for single-molecule tracking, these techniques have not yet demonstrated the observation of fast 3D dynamical processes in a cellular structure. The main limitation of the camera systems is the slow frame rate. For scanning systems, the main challenge is that fast molecules move out of the limited region for which the illumination pattern is optimized, see sup.sec.(A). However, if the reporter is moving slowly [7, 13, 22, 23] or is confined [8, 9] within a small volume, an information-optimized localization method can be used. Only a few well-placed, excitation shots are then needed to obtain a unique photon signature that pinpoints the position at high temporal and spatial resolution. For fast reporters moving in a larger volume, a more responsive and/or larger tracking volume is required. Responsive systems pose high requirements on hardware bandwidth, shot noise, and photophysics of the reporter. A large tracking volume, on the other hand, has a negative impact on the time resolution.

Here, we describe a generalized theoretical treatment of the SM-MEL principle which we also implement in a 3D single-molecule tracking system. Our single-molecule tracking implementation is based on minimizing the cross-entropy between the emitted photon counts and the true excitation PSF. It introduces a new regularization schema that stabilizes the maximum a posteriori (MAP) estimate of the position when tracking reporters moving in a larger tracking volume. This effectively reduces the bandwidth requirements of the piezo positioning system which is often limiting when tracking fast reporters. On the more practical side, we introduce a new method which removes any need for fast axial scanning equipment. In contrast to the common approach where a Tunable Acoustic Gradient (TAG) lens or fast active refocusing equipment is used for axial capability, we have introduced axial capability by encoding z information into an engineered PSF which is only moved in 2D. We test the microscope and our understanding of the system by moving and tracking fluorescently labeled 20*nm* beads and compare the results to our simulation of the tracking system. Next, we use simulations to evaluate our capability of tracking molecules in cellular geometries and inferring their binding kinetics using a diffusion point estimator in conjunction with a hidden Markov model (HMM). Finally, the microscope is unleashed on single molecules moving in live bacterial cells.

## Results

### Single molecule tracking concept

Consider the problem of estimating the unknown position 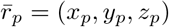 of a diffusing, fluorescently labeled molecule. The approach adopted here is based on structured illumination together with Bayesian statistics and priors which makes it possible to extend the localization problem of stationary and slow-moving reporters to more rapidly moving ones. Our labeled molecule is excited by a sequence of known point spread functions (PSF), 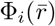 indexed by the subscript *i*, and the emitted photons, *n*_*i*_, are detected with a single-photon counting avalanche photodiode (SPC-APD). In general, the PSFs are arbitrary and include Gaussians, doughnuts, or even random speckle patterns that are experimentally measured. But, for simplicity, we assume that the same PSF, 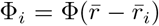, is used at the positions, 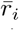. The excitation and measurement times are considered to be long compared to the fluorescence relaxation time and the emitted photons can thus be assumed to follow a Poisson distribution with a mean *λ*_*i*_. Also, we assume that the mean photon counts are proportional to the PSF intensity. Thus 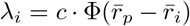 where *c* can be considered a fluorophore photon efficiency, a combination of excitation, emission, and detection efficiency per excitation power. The likelihood, given *m* PSF shots at locations 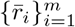, to obtain the photon counts 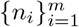from a fluorophore at 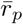 with a photon efficiency *c* can now be stated as

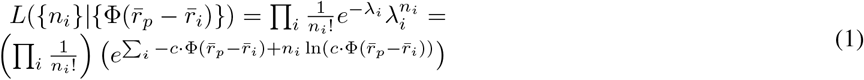

Using Bayes rule we can invert the problem to obtain the likelihood that the fluorophore is at the position *r* with photon efficiency *c* given the photon counts 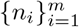 and PSF positions 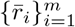. The unknown parameter *c* can be estimated by following the standard maximum likelihood procedure

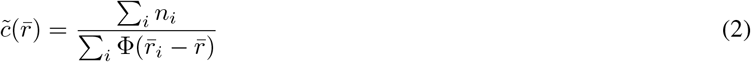

where tilde will be used to indicate estimators. The estimator is simply the ratio of the total amount of photons detected to the total PSF intensity and gives a map of the estimates of fluorophore photon efficiency over all allowed positions 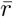. Inserting eq.(2) back into the likelihood function eq.(1) with some simplifications gives

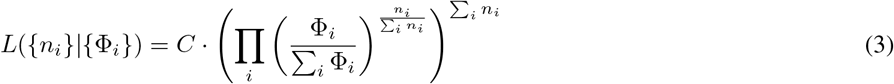

where *C*({*n*_*i*_}) is a scaling factor that depends only on {*n*_*i*_} and thus does not affect the overall shape of the likelihood landscape. It should be noted that the negative log of eq.(3) is up to a constant equal to the cross-entropy between the photon counts and the PSFs. Derivation of eq.(3) without passing through eq.(2) is also possible by the multinomial approach[7–9], but in that case, the relation to the *c* value is lost and is not accessible for construction of possible priors which are multiplicative factors to eq.(3). As described further down, constraints on the estimated fluorophore photon efficiency can be imposed, either statically or dynamically, to exclude spatial regions with very low/high excitation power compared to measured photons as described below. To emphasize the relation between the PSF ratios and the photon ratios, eq.(3) is in a form with a global exponent, Σ_*i*_ *n*_*i*_. The exponent will not change the location of the likelihood peaks and valleys but will make them sharper if increased. This is quantified by the Cramér Rao lower bound (CRLB) for eq.(3), which will depend on the total number of photons, the PSF shape, and PSF pattern used. Thus, the underlying principle of the localization by eq.(3) is to produce a series of photon count ratios {*n*_*i*_/Σ*n*_*i*_} for a given excitation pattern and find the position with corresponding ratios between PSFs which represents the expected mean photon ratios {*λ*_*i*_/ Σ *λ*_*i*_} per position. Background bias can be incorporated within the PSF model; both constant and spatially distributed bias can be added and considered a part of the PSFs. For further discussion and 1D examples, see sup.sec.(C-A).

By itself, eq.(3) needs to be confined to a relatively small region or used in a high photon count setting to produce reliable and correct position estimations. But, experiments are normally photon-count-limited and in this regime, the likelihood might have sudden large peaks that are unphysical and arise due to photon fluctuations. To counterbalance this problem, we introduce a regularization schema based on weak priors which are multiplied with eq.(3). First, a simple constraint on permitted *c* values is introduced by the binary map 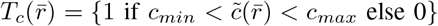 where *c*_*min*_ and *c*_*max*_ are selected depending on the PSF pattern, the PSF shape, and the sample. The upper value is the most important since it excludes regions where the sum of all PSFs is low and close to the noise floor; in these regions, the PSFs has not enough power to excite a fluorophore to obtain the registered photon counts. The second prior targets the interest of searching for positions that keep the time evolution of estimated *c* values to be on average smooth. Over a longer time span, it is assumed that the fluorophore maintains a rather stable *c* value until it bleaches. For that purpose, we assume that we can, over longer time-periods, average over the fluorophore blinking time, and apply a Gaussian with fixed width and an exponential weighted moving average as its mean. This restricts the possible *c* value dynamically over time. Parameters for both of these priors are selected to be very weak. The third and last prior utilized is to penalize large jumps within the tracking volume. This is done by applying a Gaussian with fixed widths around previously estimated positions. For example, if a width of 400*nm* is selected at 0.84*µs* full pattern update rate, this corresponds to a maximal diffusion rate of 24*µm*^2^*/s*. Further discussions and details on priors and the post-processing steps can be found in sup.sec.(C-A).

### Implementation

An overview of the optical implementation is shown in Fig.(1**a**). The excitation laser is passed through an amplitude modulator and a spatial light modulator (SLM) for PSF engineering. Two xy scanning systems are used. The first is a fast, short-range xy electro-optic deflection (EOD) system, and the second is a piezo driven tip/tilt mirror scanning system which is placed after the dichroic mirror and used for long-range xy scanning. After both scanning systems, the excitation path ends with an objective that focuses down to the sample which is on an xyz-piezo stage. Detection of fluorescence is done by a standard confocal configuration with a large 200µm pinhole and an SPC-APD. A detailed description of the optical layout can be found in sup.inf.(B-A). The SLM is programmed to produce a PSF that consists of three Gaussians, each placed on the outer rim of a circle with a 2*µm* diameter in the sample plane. One of the three Gaussians is placed on the focal plane while the other two are shifted 400nm axially in each direction. At the intermediate image plane directly after the EOD, a cropping mask corresponding to a diameter of 2*µm* in the sample plane is used to mask out the three Gaussian beams. The EOD xy-scanning system can now draw any pattern within the 2*µm* detection area on three z-planes. During tracking, a 50*kHz* pulse schema is used for excitation. The first 10*µs*, while the laser is off, is used for moving the beam into place and the remaining time is used for excitation. When shifting between z-layers, 10*µs* is added by blanking out one laser shot. The pattern used is a 13 point xy pattern repeated on each of the three z-planes plus 1 blank shot for each move between z-layers. The pattern spans a volume of 950 × 950 × 1390*nm*^3^. A representation is found in Fig.(1**b**) where each star indicates Gaussian center positions and the interconnections represent the order of the shots. The emission from each laser shot is recorded by the SPC-APDs and processed by a field-programmable gate array (FPGA). The FPGA is used to raster scan the sample, find tracking reporters, produce the tracking pattern, update the tip/tilt-piezo mirror position by calculating a centroid, as well as bundle all necessary data to be sent to the computer.

**Fig. 1.**
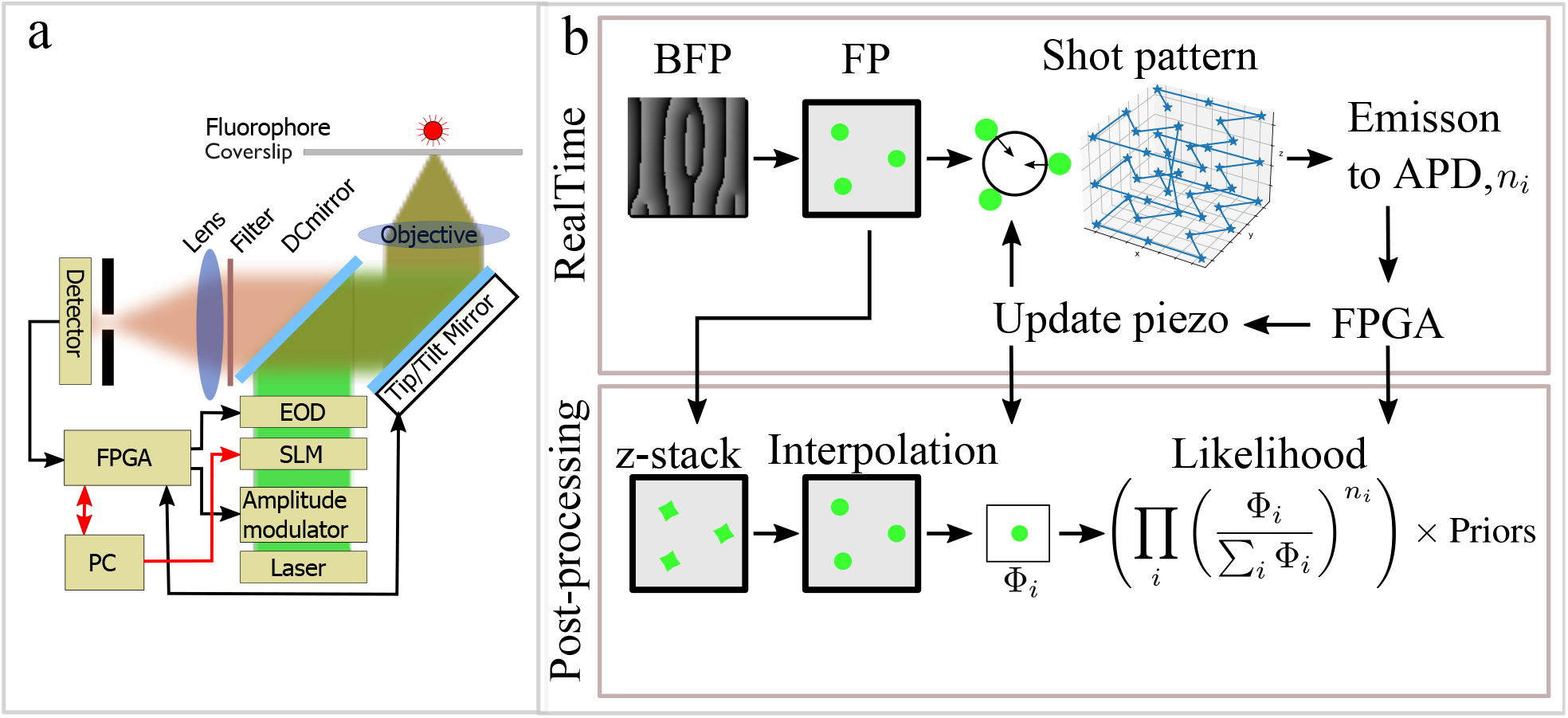
Optical setup and tracking concept. **a**, illustration of the optical setup, laser excitation path with amplitude modulation, spatial light modulator (SLM) for PSF engineering, and electro-optic deflectors (EOD). Following the excitation light path, after the dichroic mirror a piezo driven tip/tilt mirror scanner is used for long-range scanning, the light path ends with an objective and sample. Collected fluorescence is de-scanned by the tip/tilt mirror and passed through the dichroic mirror, fluorescent filters, and ends with a confocal detection based on single photon counting avalanche photodiodes. **b**, Graphical representation of the real time and data post-processing. Real-time (left to right); The PSF is shaped at the BFP so that the desired PSF appears at the focal plane (FP). The PSF is placed within the illumination area by the EOD to create the tracking pattern that excites the fluorophore. The 3D pattern of the lobes used is shown in the shot pattern image. Emitted photons are detected and processed by the field-programmable gate array (FPGA) to obtain a new piezo position. Post-processing (left to right); the PSF is sampled as a z-stack which is interpolated, and processed with the EOD pattern and piezo center. Along with the photon counts, this fully reconstructs the illumination pattern and gives the parts necessary for constructing the likelihood.

The trajectory estimation is refined in a post-processing step by maximizing the likelihood for *r*_*p*_ in eq.(3) together with its priors. To do this, we measure the true excitation PSF, Φ_*i*_, by acquiring an image z-stack using the EMCCD camera. The image stack is interpolated to obtain 10 × 10 × 10*nm*^3^ voxels; details are described in sup.inf.(B-A). Together with the EOD pattern, the tip/tilt-piezo position, and the photon counts {*n*_*i*_}, we reproduce the sequence of PSFs 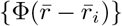 used for tracking and calculate the MAP which is our position estimation 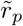.

### Experimental evaluation in combination with simulations

Evaluation of the real-time tracking system and post-processing is accomplished by tracking immobilized 20*nm* beads moved with a piezo stage. The beads are immobilized in agarose and mounted on the sample stage. The stage is programmed to move in a circle with a radius of 1*µm*, and perform a sinusoidal pattern with peak to peak amplitude of 0.5*µm* in the axial direction. Both movements have a duration of 1*s* with a subsequent 0.5*s* pause before the motion is repeated. The bead tracking is compared to simulated tracking of a fictive fluorescent particle traveling in a path similar to that of the immobilized beads, where the simulated photon counts are generated by the experimentally acquired PSF. These photon counts and PSF positions are then run through the same post-processing as the bead data. In typical experiments, the photon rate is around 60*kHz* when the fluorophore is excited with an average power of 24*µW* per PSF at the back focal plane of the objective. At this count rate, the spatial resolution is on average 67*nm* in the lateral and 109*nm* in the axial direction with a time resolution of 0.84ms (Fig.(2)**a,b**). To see how the resolution scales with the photon count rate, we apply an exponential moving average over the likelihood (eq.(3)) during reconstruction, see sup.inf.(C-A). When comparing the bead tracking with the simulations (Fig.(2)**c**), we find that the tracking error is larger for the beads compared to the simulation, although the overall trend is preserved. In general, the piezo stage appears to follow the prescribed path; however, closer inspection shows clear oscillations in the reconstructed path and that the stage overshoots its mark at the start and end of the pause. We attribute this behavior to the stage’s underdamped feedback loop as well as vibrations induced from the stage motion. These behaviors negatively affect the resolution estimation and imply that the true resolution is likely closer to the theoretical limit of 41*nm* lateral and 95*nm* axial observed in the simulation. During actual single molecule tracking, the stage is stationary in the lateral direction and only moving very slowly in the axial direction. To further mitigate any possible stage problems, we turn off the stage piezo feedback system and monitor z-axis displacements with a 980*nm* laser which is totally internally reflected at the glass water interface during measurements. This reduces any stage-induced movements and brings the resolution close to the theoretical values obtained through simulations. This means that our experimental resolution is better than what can be obtained by tracking a bead moved by the piezo stage. For further implementation details and validation tests, see sup.inf.(E-A).

**Fig. 2.**
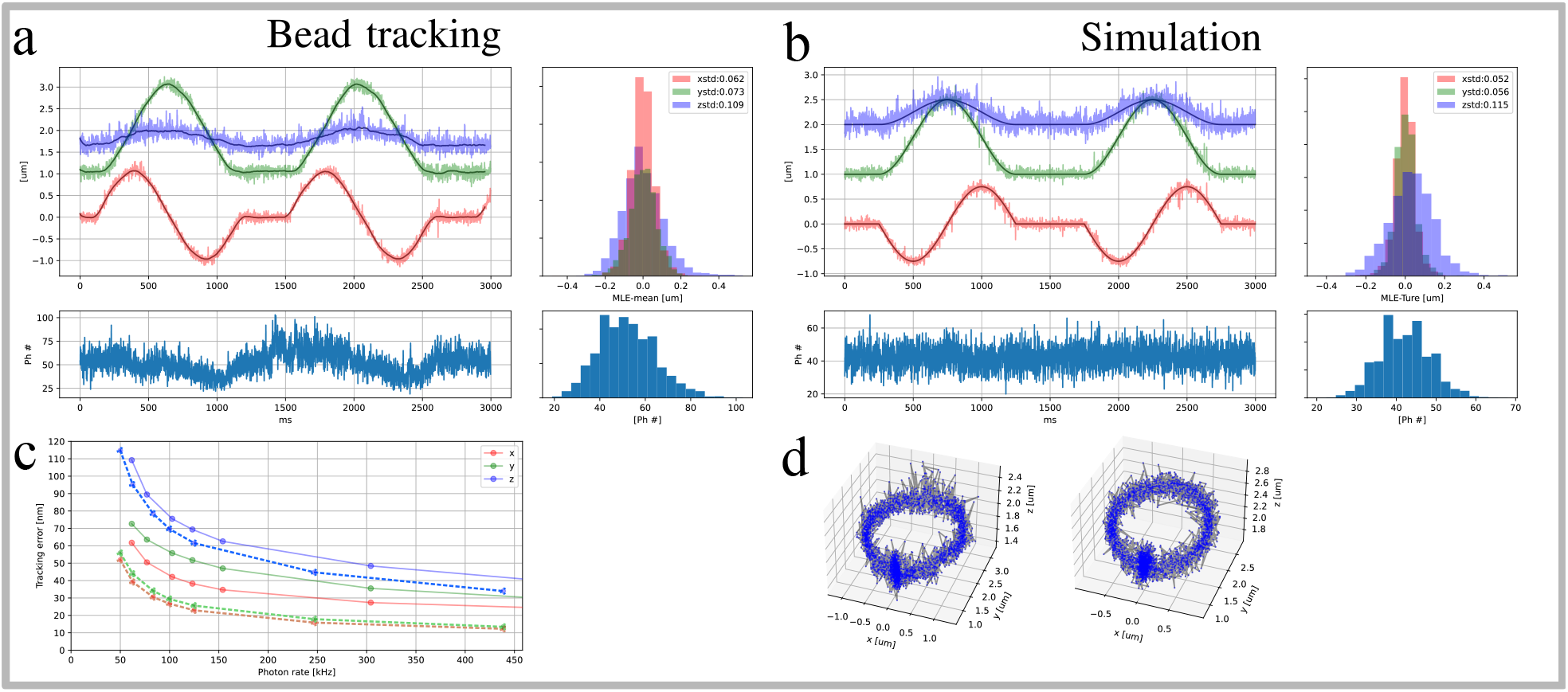
Resolution test and comparison to simulation. **a**, Tracking of beads fixed in agarose, the piezo stage is programmed to trace circles with a x,y-radius of 1*µm* and a z peak to peak oscillation of 0.5*µm*. Upper left; Trajectory estimated x (red), y (green) and z (blue) coordinates with a temporal resolution of 0.84ms. Upper right; distribution of standard deviations over the trajectory between estimation and a moving average with a window size of 100 points. Bottom left; photon counts per localization which corresponds to 60kHz photon count rate. To the right the photon count distribution. **b**, Similar to **a** but for a simulated trajectory with an x,y-radius of 0.75*µm* and a z peak to peak oscillation of 0.5*µm*, the error distribution (upper right) is here the distance to ground truth. **c**, resolution as a function of photon count rate for x (red), y (green) and z (blue), solid lines are for the bead data and dashed lines correspond to the simulations. Each point is the standard deviation of the distance histogram in **a** and **b**.These are obtained by increasing the exponential moving average of the likelihood. **d**, 3D point scatter plot of the estimated (left) and simulated (right) trajectory.

### Simulation in E. coli geometry

To get an estimation of the system’s ability to track molecules within cells, we make a diffusion simulation in a replicated *E. coli* geometry. We use a three-state model, with a fast diffusion state (6*µm*^2^*/s*) corresponding to a free protein in the cytoplasm, a slow diffusion state (0.1*µm*^2^*/s*) corresponding to long-lived binding to a larger complex, and a short-lived state with intermediate diffusion (1.4*µm*^2^*/s*) representing interrogations of possible binding sites. See sup.inf.(D) for the model parameters. The simulation generates 100 trajectories of random length between 50ms to 800ms and incorporates motion blur by updating the particle position for each new laser shot in the tracking pattern. In Fig.(3**a**), a trajectory is shown with its photon counts and its distance to the ground truth. The 3D trajectory is seen in Fig.(3**c** inset) where black is the ground truth and the yellow line is a track with 0.84*ms* time resolution. The error is larger compared to the bead tracking in Fig.(2) for the same photon count, which is mostly due to motion blur in the fast-moving parts of the trajectory. In the slow-moving parts of the trajectory, the position estimation is more accurate.

**Fig. 3.**
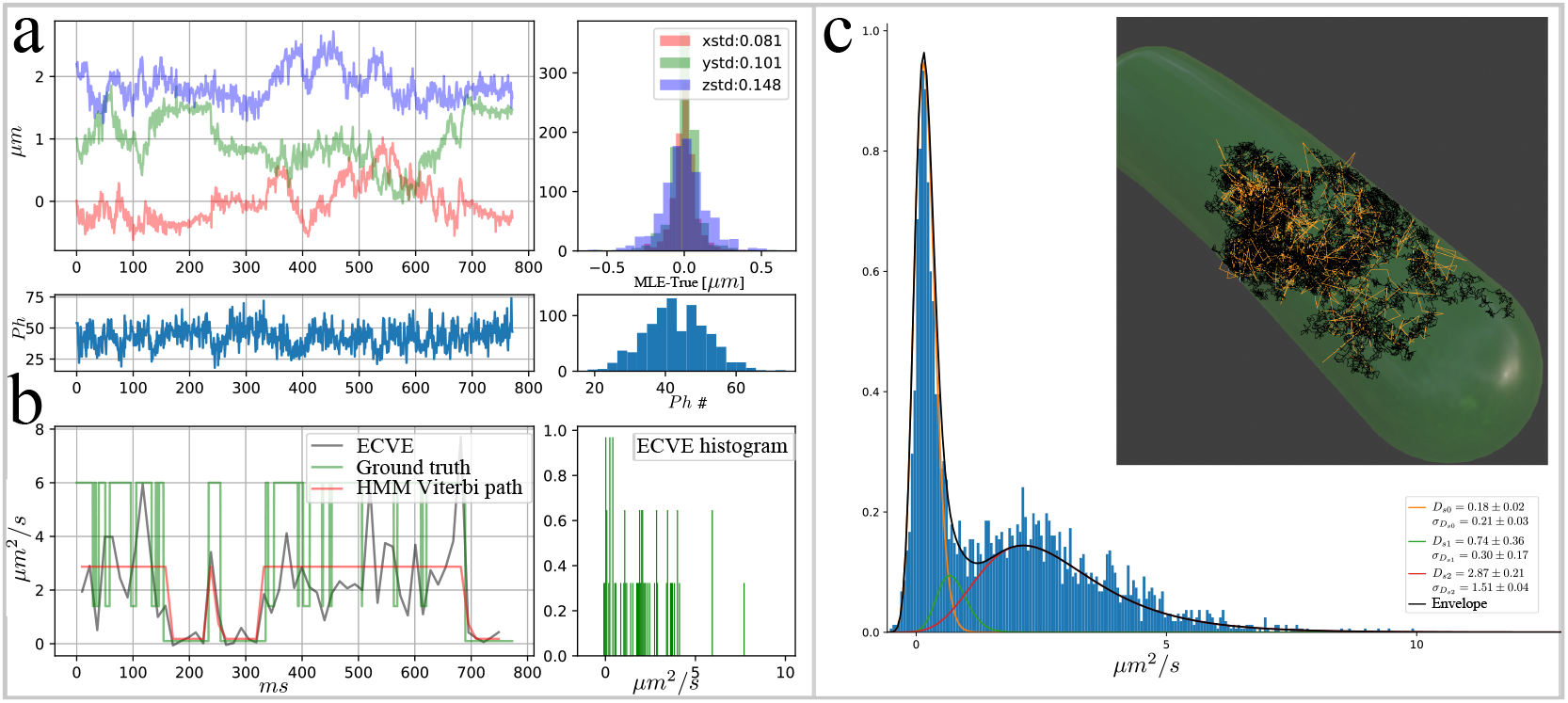
Tracking simulation in an E. coli like geometry. Example trajectory with HMM analysis of 100 simulated trajectories. **a**, upper left; example trajectory with x (red) y (green) z (blue) estimated position coordinates with corresponding photon counts (bottom). Upper right; the deviation away from ground truth with the standard deviation of the distribution in the legend, (bottom) is the photon count distribution. **b**, left; ECVE diffusion point estimation (gray) along the trajectory in **a**, ground truth diffusion is in green, and the HMM prediction in red. Right; the distribution of estimated diffusions over the trajectory in **a. c**, distribution of all ECVE values from 100 trajectories. The 3 state HMM model prediction of the emission distributions are plotted on top of the histogram, the inverse-gamma mean and standard deviation of each emission distribution is reported with s.e. from a bootstrap analysis over trajectories in the legend. Inset, 3D representation of the cell volume with ground truth trajectory in black and the estimated path from **a** in yellow.

### Evaluation of diffusion rates

To analyze each trajectory, a diffusion point estimator is derived for short intervals over the trajectory. We build on the covariance estimator (CVE) approach derived by Berglund et al. and related works [24–26], and expand their work by incorporating an arbitrary time lag (*j*) within the theory and deriving alternative point estimators. The reasoning here is that in a single step, a trajectory with high temporal resolution at low photon counts is plagued by positioning noise and will benefit from a larger time step to let the diffusion step grow while being less influenced by the localization error. By moving up in the mean square displacement (MSD) curve, larger changes are accumulated and the positioning error is less dominant. The mean square displacement over a trajectory {*X*_*k*_} is given by taking the mean over the square of steps Δ_*k,j*_ = *X*_*k*+*j*_ − *X*_*k*_ and was shown by Berglund [24] to be of the form 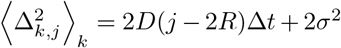when including motion blur and localization error. Here ⟨ ⟩ _*k*_ denotes the mean over the parameter *k, D* is the diffusion constant, *j* is the amount of steps, *R* is a blur factor, Δ*t* is the time of a single step, and *σ* is the positioning error. They continue and show that the covariance matrix between single steps, Δ_*k*,1_, has non-zero elements in the first off-diagonals. They use this relation to derive a point estimator for diffusion. In their treatment, the first step in the mean square displacement is given by the diagonal elements, and by taking the mean over the diagonal elements, they get the first point in the mean square displacement curve. Here we generalize their result further and obtain the covariance matrix for any time lag *j* and covariance between the steps Δ_*k,j*_. For derivation details, see sup.inf.(C-B). After some calculations, the generalized covariance matrix is given by

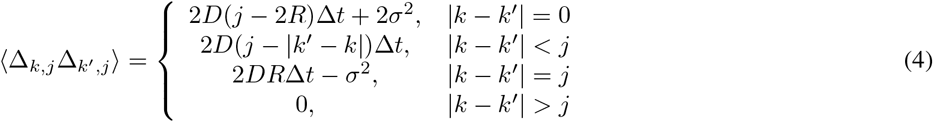

where *j >* 0 and

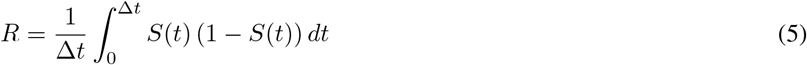

which is the blur factor with 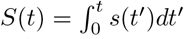 with *s*(*t*′) as the shutter function. The covariance matrix, eq.(4), tells us that moving along the mean square displacement curve (increasing *j*) indeed generates more non-zero off-diagonal elements, from Berglund’s single step (*j* = 1) with a single off-diagonal to a band of width 2*j* for a step size of *j*. It is worth noting that the localization error (*σ*) is not present for the terms in between the main diagonal and the last. Deriving a simple estimator from this covariance matrix can be done in several ways. The CVE from Berglund et al. is obtained by solving for *D* and *σ*^2^ when *j* = 1. Alternatively, but more complicated and computational heavy, one can construct a maximum likelihood estimator similar to Shuang et al. [26] but spanning over more than one *j* step. See sup.inf.(C-B) for a discussion on this topic. However, for our analysis, we note that for *j >* 1, the off-diagonals between 0 *<* |*k* − *k*′ | *< j* are all independent of *σ*, and a simple estimator for diffusion would be to solve for *D* and *σ*^2^ for a fixed value of |*k* − *k*′ |. For the first off-diagonal, |*k* − *k* | = 1, and a time lag of *j*, the estimator is

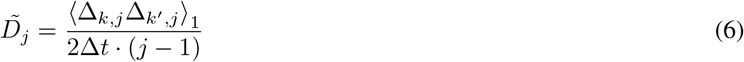

where ⟨⟩_|*k*′ − *k*|_ is the mean along the |*k*′ − *k*| = 1 off-diagonal of the covariance matrix, eq.(4). We will here refer to this estimator as the extended covariance estimator (ECVE). The localization error, *σ*, can be estimated by inserting the diffusion estimation 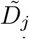 into the main diagonal, |*k*′ − *k*| = 0, and solving for *σ*^2^. The ECVE estimator is compared to the CVE and the MSD in sup.inf.(C-B1); the conclusion is that the ECVE is more robust in presence of noise and has less bias compared to the MSD when evaluated over short sections of a trajectory.

### Applying diffusion estimator to simulated data

For diffusion analysis of simulated trajectories, we use the ECVE with a time lag of *j* = 8 and averaging over 16 points. The distribution of all ECVE values is shown in Fig.(3**c**). By the shape of the histogram it is clear that it is built up by at least two distributions. The first peak seems to fit well with the slow state, the second distribution has a peak situated in between the intermediate and fast state. This is expected since the short-lived intermediate state cannot be resolved by the time steps used in the ECVE and a mixture between this and the fast state is expected. In the gray example trajectory shown in Fig.(3**b**), a clear transition between a slow and a faster state can be seen. The ground truth trajectory (green) does confirm the inability to resolve between the intermediate and fast state. It is also evident that the ECVE for the higher-diffusion parts of the trajectory is between the fast and intermediate state. Using all 100 trajectories of the *E. coli* cell simulation, we train a 3-state HMM assuming inverse-gamma distributions as emission distributions. In Fig.(3**b**), the red trace is the predicted Viterbi path, and in Fig.(3**c**), we show the distribution of all estimated diffusion values together with the predicted inverse-gamma distributions from the trained HMM. However, it is clear that the predicted Viterbi paths do not reflect the dynamics of the true interplay between the intermediate and fast states of the HMM model. Two major distributions can be identified in the histogram, the slow with mean diffusion 0.18 ± 0.02*µm*^2^*/s* (ground truth 0.1*µm*^2^*/s*), with the s.e. obtain from bootstrapping over the set of trajectories (see sub.sec(C-E)), and occupancy 48±9% (ground truth 43%) and the fast state with mean diffusion 2.87 ± 0.21*µm*^2^*/s* (ground truth 6*µm*^2^*/s*) and occupancy 45 ± 6% (ground truth 39%). The third state is identified in between the two distributions with a low 7 ± 9% (ground truth 18%) occupancy and mean diffusion 0.74 ± 0.36*µm*^2^*/s* (ground truth 1.4*µm*^2^*/s*). The short dwell time of the intermediate state is the main reason for the ECVE and HMMs difficulties to find accurate model parameters. In sup.inf.(D), we prolong the dwell time of the intermediate state and show that a more accurate HMM model fit is obtained.

#### Measurements on E. coli Trigger Factor

The methods developed above are applied to live-cell single-molecule intracellular tracking of the *E. coli* Trigger Factor (TF) chaperone system. The *E. coli* Trigger Factor is the only known ribosome-associated bacterial chaperone. One of its functions is to bind to translating ribosomes close to the peptide exit tunnel where it is believed to prevent misfolding of nascent chains during ongoing protein synthesis [27]. Previous studies have captured the binding dynamics of the TF and found a rapidly diffusing free state and a ribosome-associated state with slower diffusion due to the increase in effective size [28]. In order to track TFs in vivo, the HaloTag protein is genetically fused to the C-terminus of the TF, creating a TF-HaloTag fusion. The HaloTag covalently binds ligands, such as the organic Janelia Fluor fluorescent dyes[29], via a chloroalkane linker [30]. In the wild-type strain, the TF binds to the ribosome surface via its N-terminus. As a control, we constructed a mutant in which residues 44-46 (FRK) are exchanged to AAA which is known to result in disabled ribosome binding [31]. For details on the fused HaloTag and mutant construction, see sub.sec(H-B). Tracking experiments were performed on both the wild-type and the mutant. See sub.sec(H-C) for measurement routines. Example trajectories are shown in Fig.(4); both are selected to show the dynamical behavior of the WT and mutant TF and for that reason, they are longer than the average trajectory. In the case of the wild-type, see Fig.(4**a**), we frequently observe a bound state at low diffusion which is interrupted by occasional periods of high diffusion. All trajectories are fitted to a 3-state HMM, see Fig.(4**a**) right side where the identified HMM is plotted together with the diffusion histogram. The occupancies for the bound (0.32 ± 0.02*µm*^2^*/s*) and the intermediate (0.86 ± 0.12*µm*^2^*/s*) states are 35 ± 6% and 44 ± 7% respectively and constitute the majority of the events. Only 21 ± 3% of the time, the TF is in a more free state (2.67 ± 0.20*µm*^2^*/s*) exploring a larger volume. Looking at the mutant, we see a very different situation, see Fig.(4**b**). A bound state (0.46 ± 0.05*µm*^2^*/s*) is found by the HMM, but it has an occupancy of only 14 ± 3%. The mutant’s intermediate state (2.31 ± 0.17*µm*^2^*/s*) is here as fast as the wild-type’s free state and is occupied 35 ± 5% of the time. As seen by the diffusion histogram, there is a large tail at higher diffusion which the HMM assigns to a fast (4.81 ± 0.19*µm*^2^*/s*) state with an occupancy of 51 ± 5%. It is clear that the mutant’s average mobility is much higher than that of the wild-type, allowing the mutant TF to explore larger parts of the *E. coli* cell. Moreover, the diffusion speeds available for the mutant TF are higher than those allowed for the wild-type TF. This might indicate that what we have assumed to be the wild-type free state is in fact the TF associated with another molecule. Repeating the measurements but for a tracking pattern where the center shot is removed, leaving a hole in the pattern center, gives similar results, see sub.sec(F). No change in the resolution was observed but a notable difference in trajectory length for the wild-type, see sub.sec(G). The reson for longer trajectories is that for slow moving reporters the system reaches the MINFLUX regime at the center of the pattern.

**Fig. 4.**
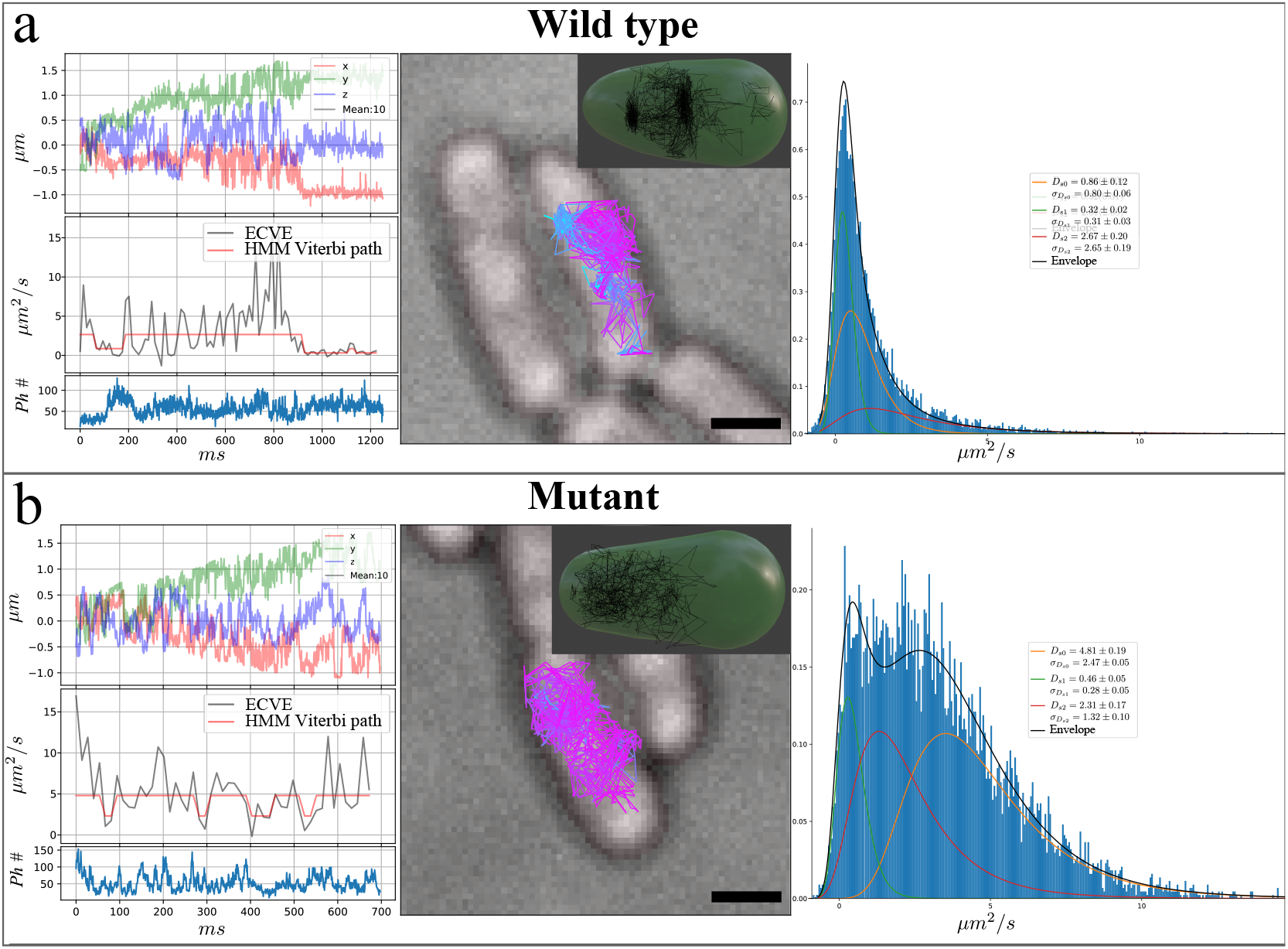
Tracking TF in WT and mutant E. Coli cells. After the microscope calibration we conducted two experiments during the same day: one experiment for the wild-type where 432 trajectories were acquired from 174 cells, and a second experiemnt for the mutant 721 trajectories were acquired from 177 cells. For each sample type, wild-type **a** and mutant **b**, an example trajectory (left) is shown together with the distribution of the accumulated diffusion histograms over all measured trajectories of each type (right side). Each horizontal block shows; Center image is the widefield cell image (black bar 1µ*m*) with estimated trajectory color coded on diffusion estimation. Inset is the 3D trajectory with a cell membrane representation based on the widefield image. Left plots, for the same trajectory shown in the center image, are from the top; position estimation, point wise diffusion estimation (gray) with HMM Viterbi path (red) and photon counts in the bottom plot. Right plot; histograms over all diffusion estimates acquired in each measurement together with the HMM emission distributions. In the histogram legend are the mean diffusion values and standard deviations with s.e. from bootstrapping.

## Discussion

We have presented a real-time 3D-tracking system capable of tracking fluorescent reporters diffusing up to 10*µm*^2^*/s* within *E. coli* bacterium. Fast 3D tracking is enabled by mitigating the workload of inertia-carrying components, like piezo-driven mirrors and stages, resulting in less physical movement when adapting to the fluorescent reporter position. There are two obvious motivations for this. Firstly, mechanical systems are slow and might induce vibrations and noise in the measurements that are difficult to separate from the tracking signal. Secondly, when tracking a diffusing molecule, the randomness in motion is not well suited for a mechanical system which carries inertia. Extending the tracking volume helps remove strain in the mechanical systems and faster molecules can be tracked. Our implementation is based on a large excitation PSF pattern and cross-entropy-based reconstruction to estimate the position of the reporter over a 950 × 950 × 1390*nm*^3^ volume achieving a resolution of at least 62 × 73 × 109*nm*^3^ at a photon count rate of 60*kHz*. For the z-axis, we adopt a novel method which removes the need of a TAG lens or other z-scanning system. Instead, we construct a set of PSFs where each PSF is at a different x,y,z location. With only 2D movement from EODs, a single PSF at a time is brought into the tracking area where a fluorescent reporter is exposed. The current speed limit of the system is mostly the update rate of the pattern (0.84*ms* for 42 shots). This is a general trade-off where systems with larger patterns have lower temporal and spatial resolution due to the extended volume they cover. Whereas systems, for slow tracking, with smaller and optimized tracking volumes can use relatively few PSF shots, and for that reason have a high positioning estimation rate. To some extent, this also has to do with the specific PSF used. Our choice of a Gaussian PSF is mostly due to convenience as Gaussians are easy to create and the simple photon count centroid estimation provides a position estimation that is easy to implement within an FPGA for piezo updates. The drawback is that the EOD system must move each Gaussian a long distance, leaving much of the volume uninterrogated for long periods of time. In our case, more than half of the time is then spent on moving the beam without interrogating the reporter. Pulsed interleaved systems may circumvent this issue but may be difficult to achieve for larger patterns. Our method together with a more optimized PSF offers an alternative. Moving away from Gaussian beams and instead using a large resolution- and speed-optimized static pattern which has a break in translational symmetry offers the possibility to shortens the time needed for beam positioning. A first attempt of pattern optimization was tested by removing the center shot of the pattern, but no obvious benefits in resolution were observed in simulations or bead tests with this pattern configuration. What was observed, when tracking the Trigger Factor with this reduced pattern, was longer trajectories for the wild-type. When the tracking pattern parks itself over a stationary reporter, we reach the MINFLUX regime which results in long trajectories. The cross-entropy principle has here been presented for the special case of positioning, but eq.(3) is not limited to PSFs and positioning. The principle is equally applicable to polarization, wavelength, or other properties where the detected photons obey Poisson statistics and can be related to a known emission/transmision profile that can be altered in a deterministic way.

Extracting valuable information from the tracking data is of equal importance as trajectory building. Molecular dynamics at the single-molecule level have been shown to be detectable by several approaches which, to mention a few, includes; multicolour tracking[32], Fluorescence correlation spectroscopy (FCS) [33], Single-molecule Förster Resonance Energy Transfer [34], and polarization enhanced FCS [13]. During our tracking experiment, polarization was measured but did not reveal any immediate signal related to conformational changes in the TF when binding to the ribosome. Detecting a correlation in conformational states and polarization signal might be possible by introducing a bifunctionally attached dye and acquiring time-tagged photon data for polarization-FCS analysis of trajectories as in [13]. Although polarization data might also carry information about the binding state of the molecule, we focused our analysis on the problem of extracting diffusion rates from positioning data on short time intervals. It was found that the MSD gives a poor estimation for this type of problem. The MSD is better suited for systems with only one diffusion rate and a small localization error. The CVE offers an alternative, but since it looks at single steps, big differences in the diffusion rates are required in combination with small localization errors. We introduced the ECVE which connects the MSD and CVE. It gives us the possibility to move up in the MSD curve, where the difference between diffusion rates is more distinguishable, and use the correlations in the covariance matrix to extract the diffusion rate with relatively few data points. An obvious drawback of analyzing displacements over longer time intervals is that we need to assume that the diffusion states have long enough dwell times; if this assumption is not valid, only an average between the diffusion rates will be seen.

In conclusion, we have reached sub-millisecond 3D tracking of rapidly moving macromolecules in bacteria, but we cannot yet resolve short-lived state transitions at this time scale. We do believe that the improvements discussed here have the potential to bridge this gap and reach the desired time and spatial resolution where macromolecular interactions occur.

## Acknowledgment

We thank I. Barkefors for helpful comments on the manuscript. This work was funded by the European Research Council (BIGGER:885360 and SMACK:947747), the Swedish Research Council (2016.06213, 2018.03958 and 2019.03714), the Knut and Alice Wallenberg Foundation (2016.0077, 2017.0291 and 2019.0439), and the Swedish National Infrastructure for Computing (SNIC) at UPPMAX.

## Author information

### Contributions

J.E. initiated the project, defined and redefined goals, E.A. conceived the method for 3D tracking using 2D beam deflection, cross-entropy minimization and the ECVE concept. E.A. and B.B. derived the theoretical framework of the ECVE and built and tested the microscope. B.B implemented the 3D PSF hologram. E.A. carried out the measurements, simulation and data analysis. T.H. developed and prepared samples of the E. coli Trigger Factor system under supervision of M.J‥ E.A., B.B. and J.E. wrote the paper, with input from all authors.

### Correspondence

Correspondence to Elias Amselem or Johan Elf

## SUPPLEMENTARY INFORMATION

Supp. info. for Real-time single-molecule 3D tracking in E. coli based on cross-entropy minimization Elias Amselem, Bo Broadwater, Tora Hävermark, Magnus Johansson and Johan Elf

## Appendix A

### 1D examples

Here we give a few examples that highlight the principles behind localization by cross-entropy minimization. Starting from eq.(3) it is worth noting that taking the natural logarithm of the likelihood and negating we get

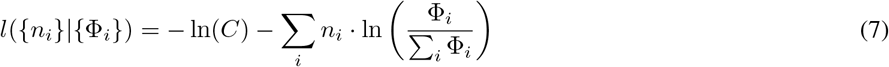

the second part can be identified as the cross-entropy between photon counts *n*_*i*_ and the PSFs 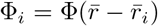 indexed by *i*, where 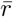 is 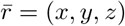 for the 3D case and 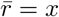 for the 1D case. The maximum resolution aloud by the model is quantified by the Cramer-Rao lower bound (CRLB) which can be stated as

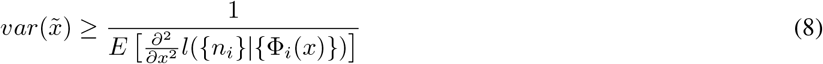

#### A. Linear case

If we consider a small region of the PSFs we can within that region linearize it to

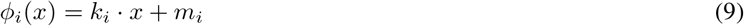

for this 1D problem consider two PSFs indexed with *i* = 1, 2. Inserting this into eq.(7) and setting the first partial derivative of *x* to zero we get the unbiased estimator

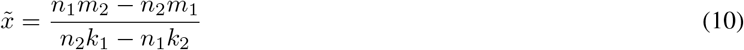

This estimator is valid in an interval where both PSFs are positive, the details of the bounds will depend on the exact *k*_*I*_ and *m*_*i*_ values. The bounds will be pairs of values given by 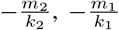or ±∞, and are independent of the photon count. Increasing the total count, *n*_1_ + *n*_2_, will result in a denser position map since there are more combinations of *n*_1_ and *n*_2_ values. As an example consider a triangular 1D-PSF as shown in Fig.(A.1). We obtain different estimators depending on which x-axis section we are looking at and how much offset is between the red and blue curve. In Fig.(A.1**a**) there are two regions, R1 and R2, and for each region an estimator is given (see the table in Fig.(A.1**a**)). The only difference between the two regions is the reversal of the sign between photon counts. Note that the estimator range is within ±*m/k* which is the range where both PSFs are positive. Also, the scaling indicates that given a fixed number of photons (*n*_1_ + *n*_2_) the possible positions will be denser with a smaller *m/k* ratio. This is the same as saying that the resolution is increasing by making *m/k* smaller, but this will sacrifice range. In contrast, increasing the total photon counts will instead increase the density of points while keeping the range.

The CRB is calculated using eq.(8) and is given by

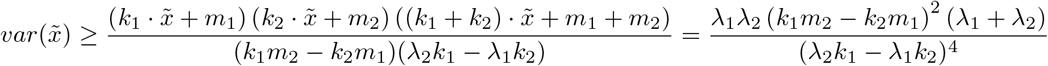

where *λ*_*i*_ are the corresponding mean photon counts. As seen in the table in Fig.(A.1**a**), the CRB (see right column in table) gives the *m/k* dependence but does not indicate anything about the range dependence. If another offset is set between the two PSFs, we get other regions and estimators. For the case in Fig.(A.1**b**), we have three regions. The first region (R1) is as in Fig.(A.1**a**) but with a shorter range since the *m* value has decreased and is bounded in the interval ±*m/k*. The second region (R2) has two parallel PSF intensity curves. The estimator gives values from *m/k* to ∞ which is valid if these continue to increase, but as seen in the plot the valid range is from *m/k* to *c*. The last term in the CRB for R2 shows that the resolution drops further away from the point *m/k* we get. The third region (R3) is as the first but with an intensity offset, this offset increases as *c* increases. The estimator has the possible range of *m/k* to 2*c* + *m/k*, but for our example it is only valid within *c* to *c* + 2*m/k*. From the CRB for the R3 region we do see that the resolution goes down as *c* increases.

Thus, to optimize the resolution, given fixed mean photon counts, one can either increase the slope while keeping the intensity bias fixed or have the PSFs intersect close to the low amplitude part and keep the slope fixed. The first option depends on the optical hardware used and has often limited possibilities for improvement. Some optimization can be done by selecting an appropriate PSF and pattern; however for most applications Gaussian or doughnut beams are preferred, and the corresponding optics are configured optimally for highest possible resolution. The second possibility for resolution optimization is the *m* value, here the limitations are related to background noise, PSF low intensity lobes, and excitation power. Our example in Fig.(A.1) is not realistic, in reality there are optical bandwidth limitations which couple the PSF *m* and *k* values. For example, a zero mean Gaussian beam has an *m/k* ratio scaling as 1*/x* indicating an increase of resolution by moving the Gaussian PSFs apart from each other. In experiments, true Gaussians are not possible. Instead the PSF has lobes which will limit resolution by separation. For a doughnut beams, there are two regions: one close to the center that has a ratio scaling as *x* and a second region, which is the outer part of the doughnut, that scales as the Gaussian. Thus, for the doughnut center the resolution decreases as the displacement is increased.

**Fig. A.1.**
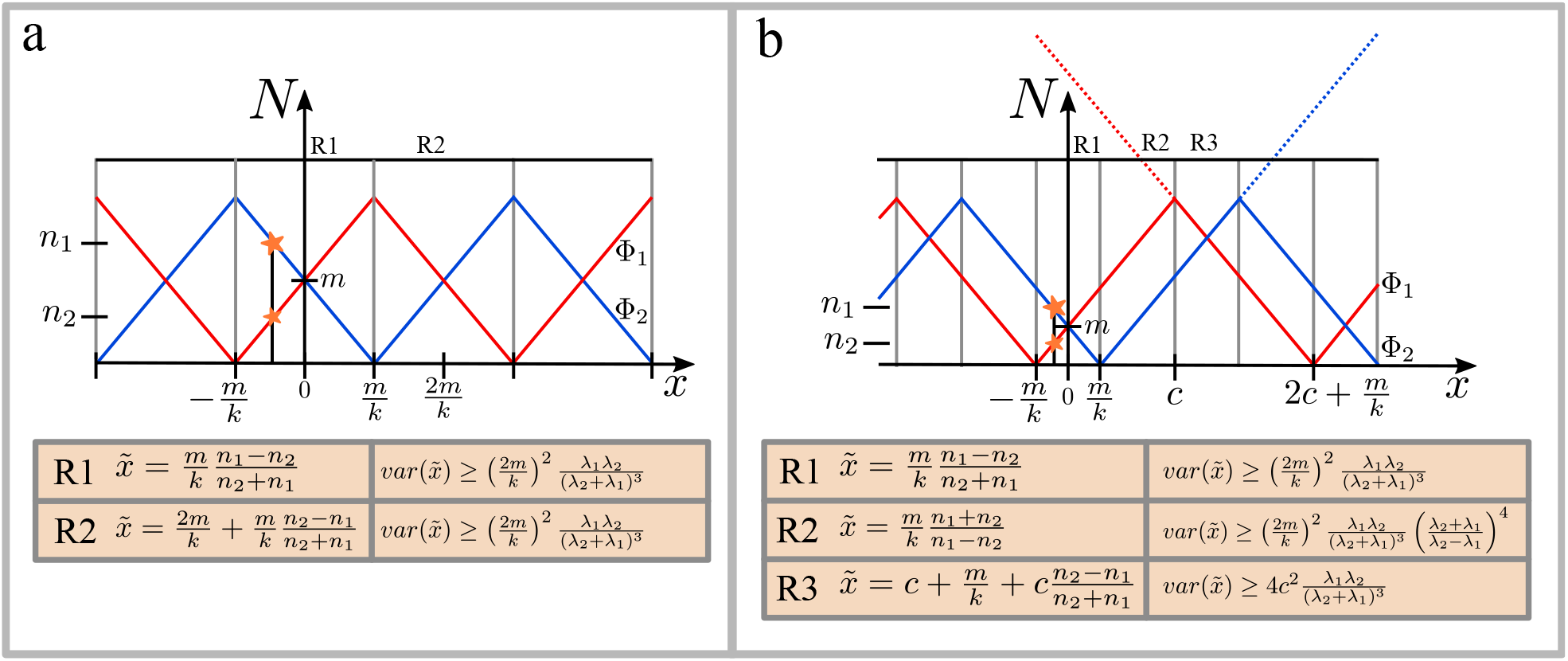
**a**. Plot shows two triangle 1D-PSFs, red and blue, over the x-axis with the y-axis being their respective intensity. A fluorophore at a position on the x-axis will emit photons proportional to each PSF intensity (marked as stars). Regions where estimators are valid are marked with gray vertical lines. In the table under the plot are the estimators for the regions R1 and R2. The first column is the estimator and the second is the CRB. **b**. the same as **a** but the two PSF have a different offset between each other. Three regions are identified, R1, R2 and R3. For each region the estimator is shown in the table under the plot. Note that each estimator is only valid within the interval marked by the gray lines in the plot and are given by the interval *±m/k* for R1, *m/k* to *c* for R2 and the R3 estimator is valid within the interval *c* to *c* + 2*m/k*. Still the estimator itself is not bounded by the interval. If photon shot noise is introduced when using the R2 region the estimator might give a wrong position since the estimator has a range from *m/k* to *∞*.

#### B. Gaussian

As a second example we describe the special case of a 1D Gaussian PSF. Consider a PSF described by

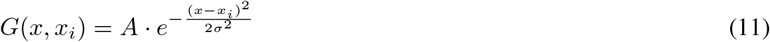

where *x*_*i*_ is the offset, *σ* is the PSF standard deviation, and *A* is related to the beam intensity. For this 1D case, we need at least two beams for a localization, *i* ∈ [1, 2], and without loss of generality, we can assume that these are placed symmetrically around the origin, *x*_1,2_ = ±*α*. Inserting our assumptions into eq.(3) gives the likelihood function

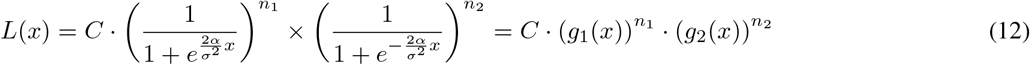

In Fig.(A.2) we plot *g*_1_(*x*) and *g*_2_(*x*) for different values of 2*α/σ*^2^, as seen the slope increases as 2*α/σ*^2^ increases. This indicates an increased resolution but only at the center, further out the resolution is decreasing.

Assuming that there is a fluorophore at position *x*_*p*_ generating photon counts, we have the problem *L*(*x*_*p*_|{*n*_*i*_}). For a general PSF together with a given pattern there might be several maxima, thus finding a unique maximum likelihood estimator might not be possible without further assumptions. However, for the Gaussian example, there is a unique solution. Following the standard maximum likelihood procedure gives

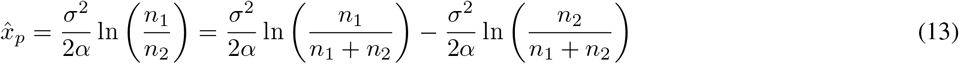

which is our unbiased estimator. The localization performance can be calculated from eq.(8) and eq.(12) which gives

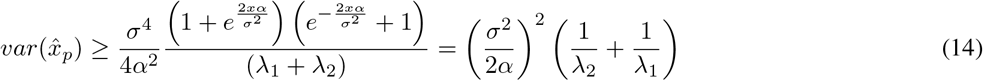

where *λ*_*i*_ are the mean photon counts. As can be seen the CRB is not uniform over *x*, as we move further out from the center we gradually lose resolution. As expected the resolution decreases when the mean photon rate decreases. As long as one is at the center, it is more beneficial to increase the distance, 2*α*, between the beams or decreasing *σ* compared to increasing the photon count by higher excitation power. This can also be seen from eq.(13) where for any non zero, finite total photon number, *n*_1_ + *n*_2_, the estimator covers all space, with the limits ±∞ when *n*_1_ or *n*_2_ are zero. But the total number of possible points between ±∞ will be finite, more precisely *n*_1_ + *n*_2_ − 1, and these will span an interval on the x axis of length (*σ*^2^*/α*) ln (*n*_1_ + *n*_2_ − 1). Thus increasing the distance between the beams will result in a higher sampling density at the center. But this effectively decreases the interval when we exclude extreme points at ±∞. In contrast, increasing the total photon count will add points over the whole interval. Thus, for imaging applications where only slow movements of the reporter are present a resolution optimization using the beam configuration can be done by sacrificing range. But, if the reporter is moving one needs to increase the range of the estimator by sacrificing resolution.

**Fig. A.2.**
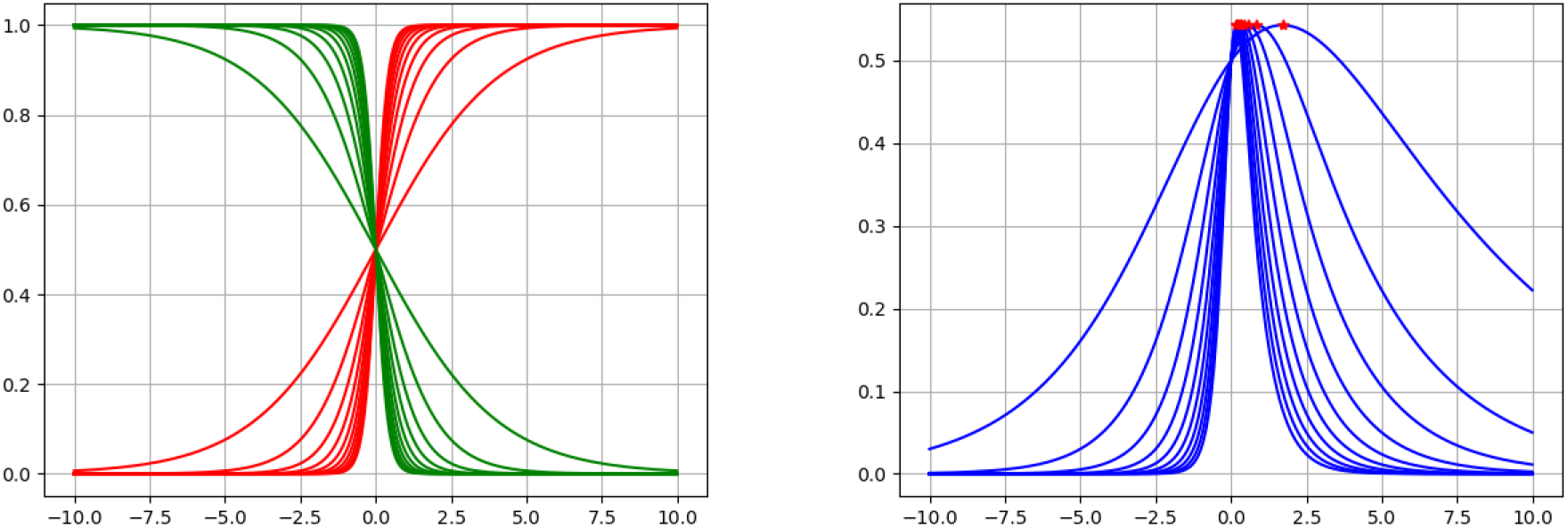
Left plot shows curves for the *g*_*i*_(*x*) part of eq.(12) with 2*α*/*σ*^2^ = *k, k* = [1,‥,10]/2. As *k* increases the steeper the function transits from 0 to 1. Red is *g*_1_(*x*) and green is *g*_2_(*x*). Right plot is 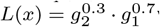which represent the likelihood over the positions x for different *k* values. The exponent corresponds to the percentage of the mean photon counts, λ_*i*_, with 0.3 = λ_2_/(λ_1_ + λ_2_) and 0.7 = λ_1_/(λ_1_ + λ_2_). As seen the likelihood is more narrow for high *k* values. The red points indicate the maximum, they do not coincide since for different *k* values the photon counts used represent different positions.

## Appendix B

### Optics and hardware implementation details

In these subsections, we present the hardware used and related optical methods as well as the implemented software. The scanning/tracking microscope software implementation is custom built and based on Labview and Labview FPGA. The EMCCD camera was operated under MicroManager and the focus tracking system was controlled with a custom Python script.

#### A. Optical implementation

A detailed view of the real-time tracking microscope is shown in Fig.(B.1). The working principle of the microscope is to first have an amplitude modulator based on a Pockel-cell, which is followed by a TEM00 mode cleaner implemented by a pinhole. From here the beam is expanded and projected on an SLM where we tailor the PSF of the system. Following the beam, we enter into the first x/y-scanning stage based on EODs. After exiting the first scanning stage, we clean the polarization and rotate the light by half and quarter wave plates. This brings us to the dichroic mirror which is directly followed by the second scanning stage, a tip/tilt piezo system. From here we bring the laser light to the objective and the sample piezo stage. Collected fluorescence is de-scanned by the second scanning system and transmitted through the dichroic mirror and focused through a pinhole, light is then brought to a wollaston prism and finally to the SPC-APDs.

For widefield imaging the excitation light is coupled to a multimode fiber and at exit a rotating diffuser is used before entering the microscope through a flip mirror. An EMCCD camera is used for alignment and viewing samples.

Monitoring the sample axial position was done by a 980nm laser that is passed through the objective in totally internally reflected mode, the reflected light was collected by a CMOS camera that is triggered each 7.56*ms* by the FPGA when tracking.

Equipment used is found in the following list:

- Laser: Coherent Verdi 532nm
- Pockel cell: New Focus 4102NF
- SLM: Hamamatsu LCOS-SLM X10468
- EOD: Conoptics 311A
- Tip/tilt mirror: PiezoSystem jena PSH 10/2 SG with d-drive
- SPC-APD: Excelitas SPCM-AQRH
- EMCCD: Andor iXon Ultra 897
- Objective: Nikon Apo Lambda 100x 1.45Na
- Stage: Thorlabs NanoMax stage
- Dichroic mirror: Semrock FF552-Di02
- Emission filter: Semrock FF01-585-40
- Excitation filter,:Semrock FF01-770/SP-25 and FF01-532-18
- Pinhole, 200*µm*Thorlabs
- CMOS camera: Allied Vision Stingray
- Lenses: Thorlabs Achromatic Doublets

**Fig. B.1.**
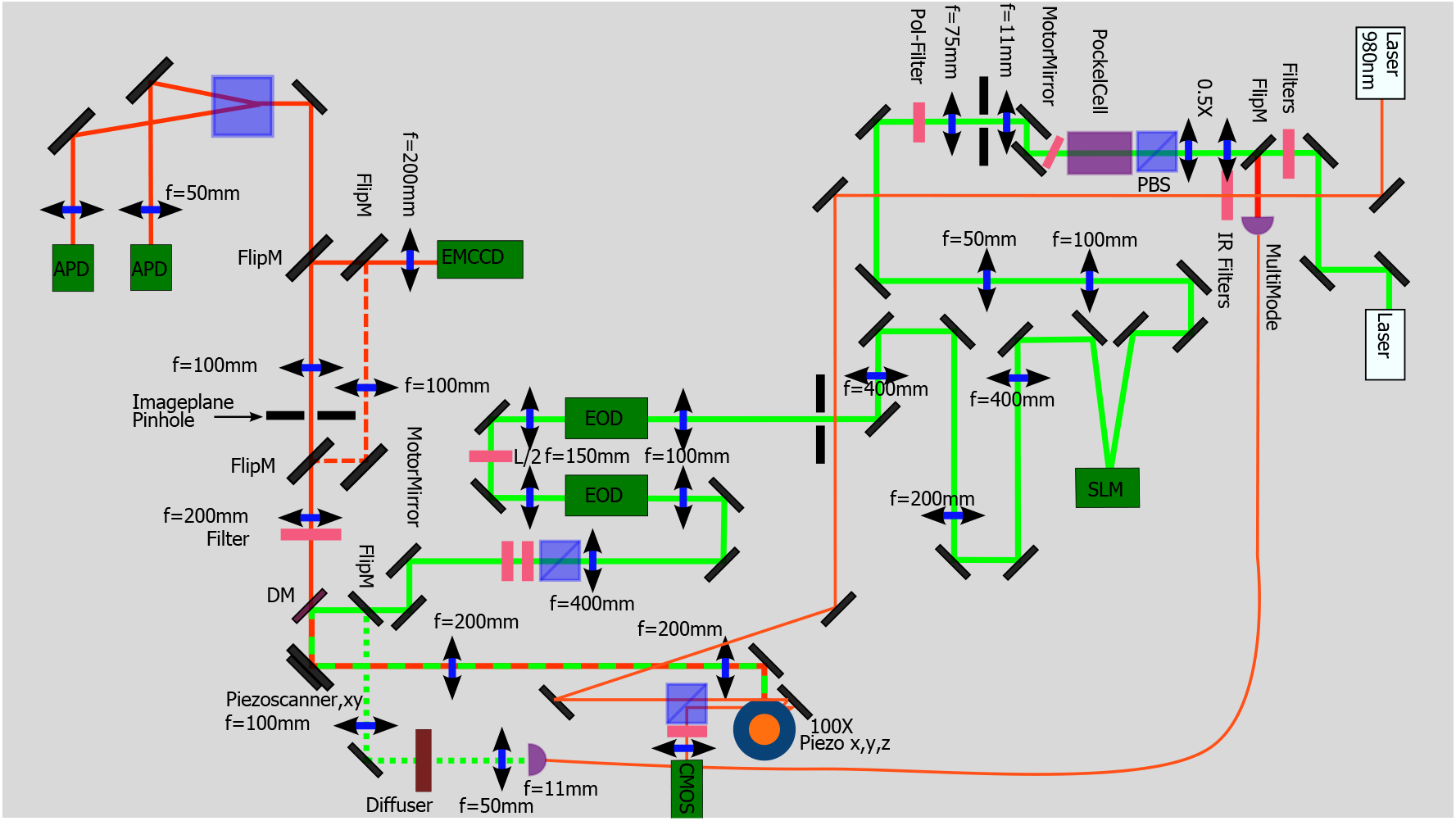
Detailed view of optics. Green beam path represents the excitation and the red path is the emission path. Orange path represents the total internal reflection focus system.

#### B. PSF engineering

We engineered the PSF for the excitation light with a spatial light modulator (SLM). The laser is expanded to fill the SLM where a computer-generated hologram (CGH) is drawn. Using the Weighted Gerchberg Saxton algorithm [35] in combination with aberration corrections patterns, we create three Gaussians at different axial depths.

#### C. PSF sampling and processing

The z-stack of the engineered excitation PSF is obtained by a reflecting sample placed on the piezo stage. Further, by replacing the emission filter with an OD1 natural density filter, one can observe the laser reflection onto the EMCCD camera. To test that the dichroic mirror only attenuates the laser and not alter the PSF a 50/50 beam splitter and camera was placed between the dichroic and objective to sample the PSF. This confirmed that excitation light sampling can be done through the dichroic mirror. The z-stack was sampled by moving the piezo stage in steps of 25*nm* through the PSF (50 steps), this gives a range of 2.5*µm* with 50*nm* steps and the in-plane camera resolution is 80*nm*. The z-stack is interpolated to obtain 10*nm* voxels and cropped to volume of suitable size. Interpolation is done in two steps, first in z and then in the xy-plane.

##### 1. z-interpolation

To interpolate over the z-axis, a phase retrieval is done by a double weighted Gerchberg Saxton algorithm [36, 37] which iterates between each pair of adjacent images in the z-stack. After phase retrieval the z-stack represents a light field stack with phase information for each z-stack image. By forward and backward propagation of the light field between two adjacent z-planes and taking the linear interpolation average of the two light field intensities one obtains any z-plane in between.

##### 2. xy-interpolation

After z-interpolation each z-layer is xy-interpolated by the OpenCV python package resize using the bicubic interpolation over 4 × 4 pixel neighborhood.

#### D. Real-time tracking implementation

The real-time tracking system is implemented on an National Instruments PCIe-7852R FPGA. The timing schematic is shown in Fig.(B.2). Three loops are used; first is the 50*kHz* loop that controls the piezo and EOD position, this loop is switching between scanning mode and tracking. Second, is a photon time tagging loop running at 200*MHz*. This loop controls the laser intensity, delays between laser on/off states and the photon measurement on/off states, sampling the piezo stage position, photon time tagging and photon accumulation over the measurement window. The third loop calculates a centroid estimation based on a sliding window over two full tracking patterns. This calculation gives the distance to the tracking pattern center and is used in a PID controller which produces a compensation signal to keep the reporting particle within the tracking region. The PID signal is sent to the first loop which will update the piezo position. PID parameters are tuned by hand while tracking beads and monitoring the sampled piezo stage signal to avoid oscillations. When the system is not tracking, a raster scanning is performed. Whlie scanning, the system searches for a tracking candidate. The switch to tracking mode is triggered by a photon count threshold.

**Fig. B.2.**
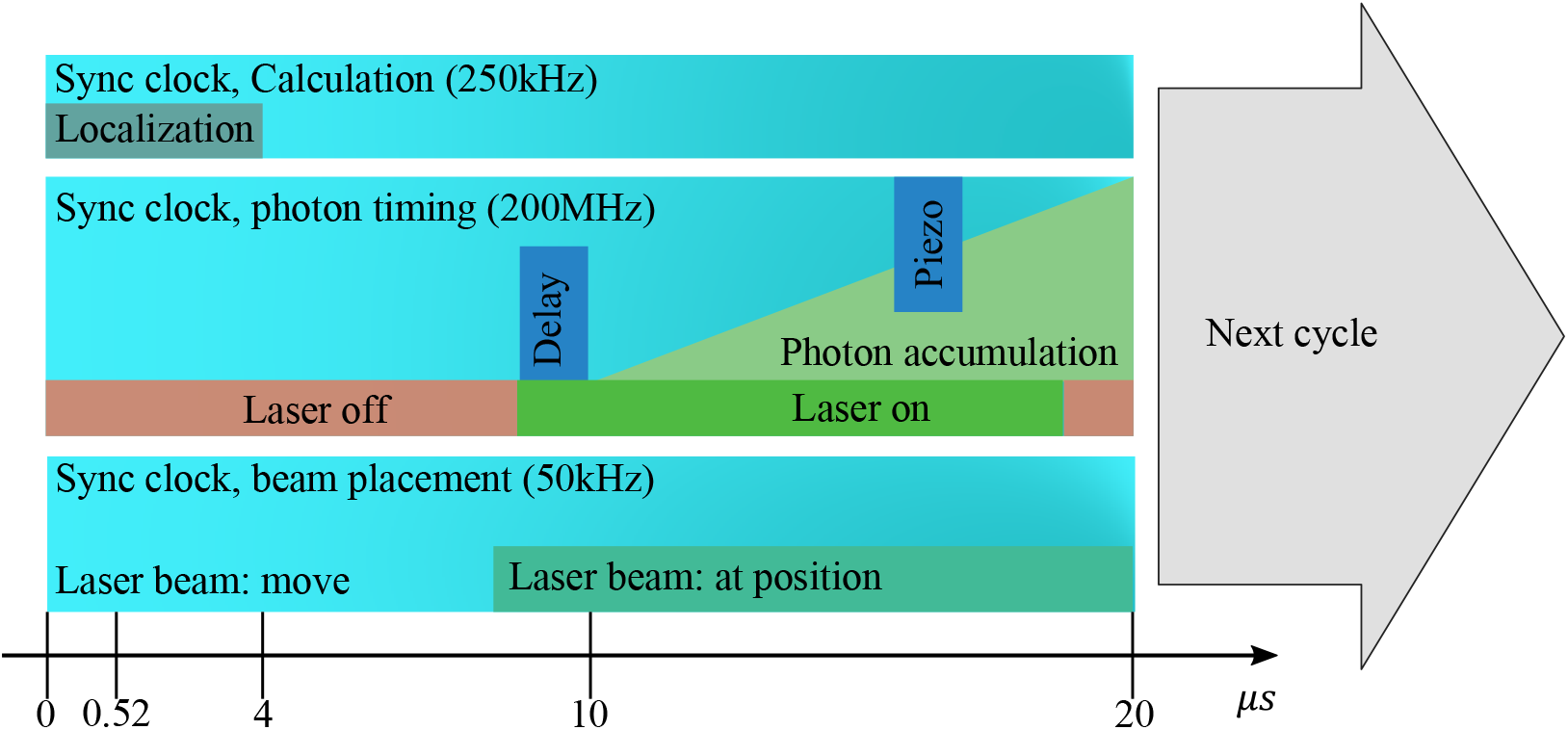
FPGA timing configuration. This illustrates one measurement cycle. The upper loop calculates a centroid estimation and also bundles the data to be sent to the computer. The middle loop handles timing of the laser on/off, measurement delay, photon accumulation and sampling of piezo stage position. The final loop initiates each new beam movement and triggers the middle loop.

## Appendix C

### Post-processing analysis details

Discussed here are the regularization methods applied to get an estimate suited for moving particles. We then introduce the point estimator for diffusion.

#### A. Position estimation

During a measurement run, we obtain a set of photon counts,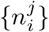,where an element 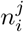 is indexed by *i* ∈ [1, ‥, *m*] corresponding to the PSF 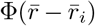 that builds up the full pattern, and *j* ∈ [0, ‥, *M* − 1] which are the amount of full patterns acquired during tracking. One full pattern, see main fig(1), consists of *m* PSF shots. In our implementation *m* = 42 including the three blank shots which are defined to be zero for both photons counts and PSF, thus these do not contribute to the likelihood. Using the PSF in sec.(B-C) together with the tip/tilt mirror piezo position and the EOD position we obtain an the experimental measured PSF 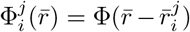 with known 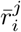 and discretized 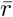 over a grid of size 950 × 950 × 1390*nm* with 10*nm* voxels.

The likelihood (main eq.(3)) can be restated as the negative log likelihood, and with the indexing convention introduced above, is transformed to

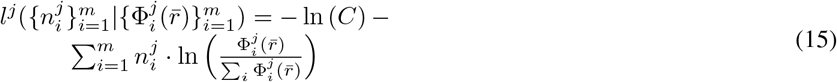

here the second part can be identified as the cross-entropy. When tracking slow moving reporters at high signal-to-background ratio with a well behaving PSF in a small search volume, one can seek a position estimation directly from eq.(15) by 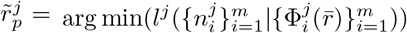 over possible positions 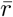. However, it is necessary to regularize eq.(15) for larger search volumes when the signal-to-background ratio is decreased and/or the tracking reporter is faster. One method is to use weak priors. Here, we use time dependent physics motivated priors to regularize possible estimations. We seek an estimator of the form

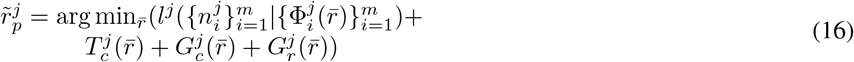

where each prior is defined in the following subsections.

##### 1. Prior 1

The fluorophore photon efficiency, main eq.(2), is bounded to exclude regions that fall outside the expected range. This effectively removes regions where the search volume has a total intensity that is too low to be considered a region where any significant photons signal originate. The upper bound is set high enough to not affect the likelihood. Thus, this binary map, after taking the logarithm, is given by

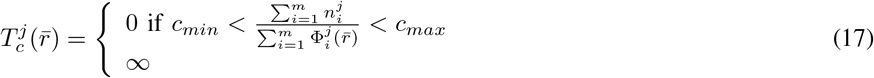

where *c*_*min*_ and *c*_*max*_ are selected depending on the pattern and PSF shape used. In practice 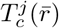 is independent of *j*, but there might be situations where the PSF pattern is either more dynamic and the binary maps are not the same between *j*’s or that bounds need to change depending on the tracking situation. Note that the lower bound restricts the lower total intensity allowed. In the case of optimizing for photon flux, as in MINFLUX, one might exclude important regions.

##### 2. Prior 2

The previous prior only gives the outer bounds of possible fluorophore photon efficiency values. During position estimation a more dynamic but tighter constraint is of interest. Over a longer time span, it is assumed that the mean photon counts per excitation power is not changing widely but is changing rather smoothly. To incorporate this as a weak prior we make an exponentially weighted mean of *c* values obtained from estimated positions 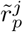, this defined in a recursive way by

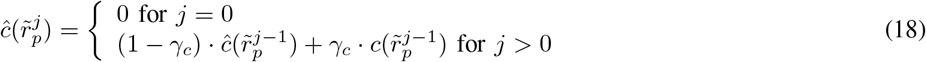

where *γ*_*c*_ is the weight parameter, and is set to have a long tail that will average over blinking events. With this averaging, a gaussian prior is constructed and after taking the logarithm gives

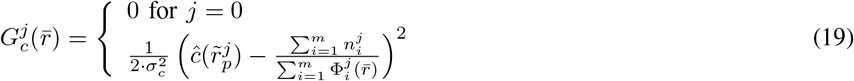

where *σ*_*c*_ is a fixed parameter defining the allowed span of the gaussian.

##### 3. Prior 3

The last prior is a gaussian prior on the previous position estimation, its effect is to suppress very long jumps assuming that the next position is probably closer to the previous estimated positions then far away. Here also we use an exponentially weighted mean but on the estimated positions

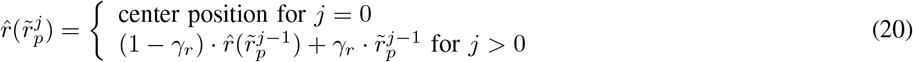

where *γ*_*r*_ is the weight parameter, and is here chosen to have a short tail. The prior is then given by

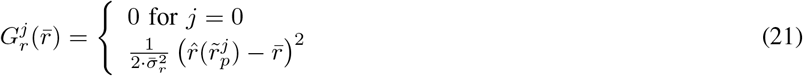

where 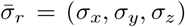 is a fixed vector of values defining the reach of the gaussian. The value is set depending on the sampling speed in relation to expected max diffusion of the sample to only have a weak effect.

##### 4. Likelihood exponential averaging

Increasing the resolution can be done by increasing photon counts. In post-processing we can sacrifice timing bandwidth assuming that the reporter is moving slowly. A way to indirectly do this is to take the mean of estimated positions to get a refined position. Or one can sum photon counts emanating from the several PSF shots at a position and estimate positions. Here, we instead construct an exponential averaging of the likelihood before adding any priors. The benefit of doing this in this way is that one is updating the current likelihood by taking into account that current likelihood is the most relevant but that the history of past likelihoods should be considered in trailing fashion. For this we construct a likelihood tail as

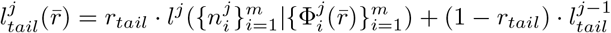

where *r*_*tail*_ is how much tail that should be considered, if *r*_*tail*_ = 1 then we are only considering current data without any historical considerations. The priors are only added after the tailing, thus the final position estimation is

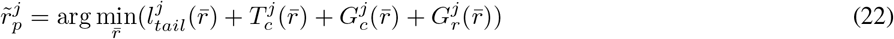

This averaging is only applied when testing resolution for beads and simulations, for all TF data we use *r*_*tail*_ = 1.

##### 5. Determining Diffusion Coefficients

Given position estimations for a particle trajectory, we like to find relevant parameters that characterize the underlying dynamics related to it. For dynamical processes where the molecule in question is driven by a diffusion process and associates/dissociates to a larger molecule, it is expected that the diffusion state-change can be observed. A more careful treatment of MSD where contributions from motion blur and position estimation noise are incorporated reveals a richer theory with alternative estimators. Building on the covariance point estimator (CVE) approach, that was derived by Berglund et al. and related work [24–26], we expand their work further to incorporate an arbitrary time lag within the theory and derive alternative point estimators.

Suppose that a particle is moving in one dimension by pure Brownian motion with a diffusion coefficient *D* and that the particle position *X*_*k*_ for *k* = 1, …, *N* is estimated with a time interval of Δ*t*, which will be referred to as the frame time, and *k* is the frame index. For higher dimensions, 2D and 3D motion, we are assuming that each dimension is independent and each can be treated separately. The observed position of the particle is then the average of its position weighted by a “shutter function” *s*(*t*), a non-negative function whose integral over the frame time interval is unity. On top of this, we add a noise factor representing the localization noise term which we assume is a zero mean gaussian with covariance ⟨*ϵ*_*i*_*ϵ*_*j*_⟩ = *σ*^2^*δ*_*i,j*_. Given that we have *N* frames, the kth frame ending at time *t* = *k* · Δ*t* where *k* = 1, …, *N*, gives an observed position *X*_*k*_ given by

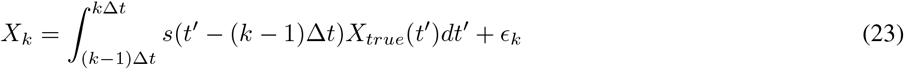

where *X*_*true*_(*t*′) is the true position of the particle at time *t*′ and *ϵ*_*k*_ is the value of the additive localization noise in frame *k*. In the ideal case when *ϵ*_*k*_ = 0 and *s*(*t*) is a delta function then the observed position is the true position and the distribution of position intervals Δ_*k,j*_ = *X*_*k+j*_ − *X*_*k*_ for any fixed *j >* 0 are independent and are zero mean Gaussian distributed with a variance 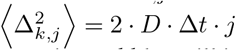 and the covariance matrix off-diagonal elements 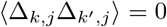 for all *k* ≠ *k*′. Introducing localization noise and blur will induce correlations between Δ_*k,j*_ which will give a covariance matrix with nonzero off-diagonal elements.

###### a) Derivation of the zero mean

Using eq.(23) it is straightforward to show that ⟨Δ_*k,j*_⟩ = 0.

###### b) Derivation of covariance

For the covariance we like to evaluate ⟨Δ_*k,j*_Δ_*k*′_, _*j*_⟩, and after some algebra and using the Brownian motion property 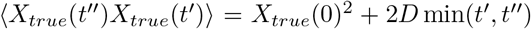 and that min(*x, y*) = (*x* + *y*)*/*2 − |*x* − *y*|*/*2 we get the covariance matrix

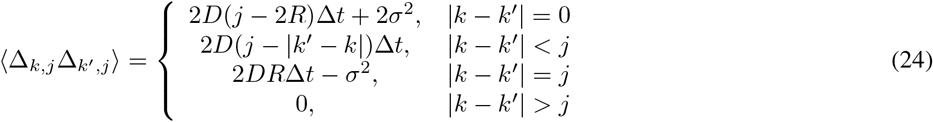

where *j >* 0 and

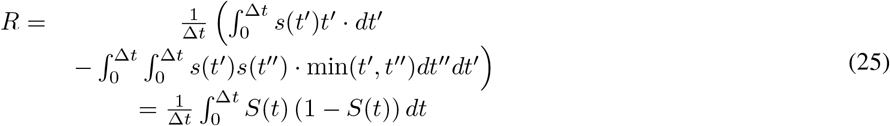

which is the blur factor with the cumulative shutter function

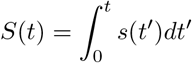

The covariance matrix diagonal elements are now the MSD curve as a function of time lag when viewed as a function of *j*. The covariance matrix presented in Berglund et al. [24] is recovered when *j* = 1, and it can be seen from eq.(24) that it is a real symmetric band matrix where the band width increases as *j* increases. Following the CVE method outlined in [38], we solve for *D* and *σ*^2^. There are several possibilities for this and we look at three:

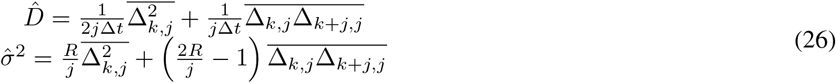

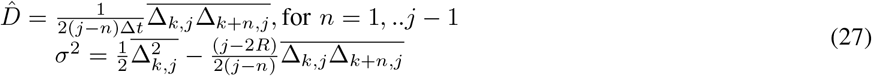

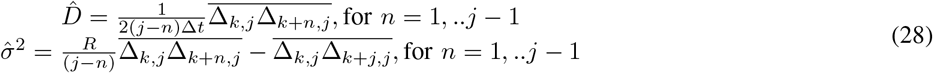

here hat refers to estimations and the bar is the mean taken along the diagonal of the covariance matrix, thus 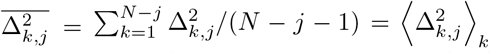, and 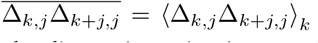 where we use ⟨⟩_*k*_ to indicate the mean over the parameter *k*. The first pair is constructed by the diagonal matrix elements for |*k* − *k*′| = 0 and |*k* − *k*′| = *j*. This is a direct extension of the CVE [38], and are the same when *j* = 1. For the second and third equations, we note that for 0 *<* |*k* − *k*′| *< j* there is no *σ*^2^ or *R* dependence and thus *D* can be estimated directly. For *σ*^2^ there are two choices given this *D* estimator. In the second pair we use the |*k* − *k*′| = 0 elements, and in the third pair we use |*k* − *k*′| = *j* elements. The estimator finally used is the second pair with *j* = 8 and |*k* − *k*′| = 1.

###### c) Likelihood function

It is possible to find an inverse to the covariance matrix, eq(24), [39], and by the spectral theorem a real symmetric matrix is diagonalizable by orthogonal matrices. Thus we can argue, as in [26], that there is a orthogonal matrix ***P*** such that ***P* Σ*P*** ^*−*1^ = ***Λ*** where 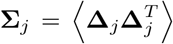 is the covariance matrix, eq(24), where **Δ**_*j*_ is the column vector with displacement of distance *j* and ***Λ*** is a diagonal matrix with diagonal elements described by the vector **λ**. Note that this also implies that there is a basis where the measurements **Δ**_*j*_ are decoupled, since the diagonal matrix 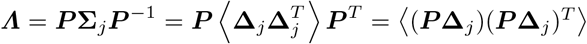. In this basis the measurements ***P* Δ**_*j*_ are decoupled and constitute a Brownian motion with step sizes **λ**. Thus, the probability for each step is given by a Gaussian, and the total log likelihood is the log of the product of all of them. Skipping all constant terms, we get for a given offset *j*

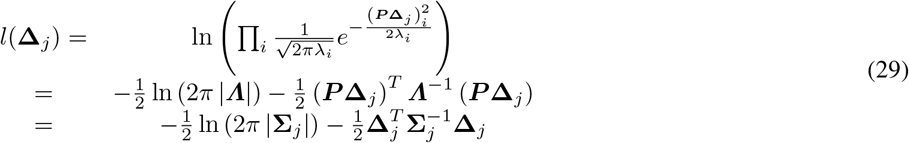

To include several *j* offsets in the likelihood, one can consider an interval of *j* values to use in the likelihood, 1 ≤ *j* ≤ *J* wich gives

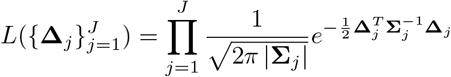

Since **Σ**_*j*_ depends on *D*, a maximum likelihood estimator can be obtained by finding *D* that maximizes the likelihood above. Since this is not a simple estimator, we will not pursue this and instead use the estimator in the main text.

###### 1) Comparison between MSD, CVE and ECVE diffusion estimations

Above we discussed the ECVE method and its relation to the mean square displacement (MSD) and the covariance estimator (CVE). To test their performance, we simulate simple 3D diffusion where the diffusion rate is increased in steps (0.1*µm*^2^*/s*, 1.4*µm*^2^*/s* and 6*µm*^2^*/s*) where each step is 168*ms* long. The time step used in the simulation is 0.84*ms* with a subsampling of 10 samples per time step. On top of the 3D trajectory, we add Gaussian noise with a standard deviation of 70*µm* per axis. Each axis of the trajectory is shown in Fig.(C.1) where the coloured section corresponds to a diffusion rates. Three point estimators are tested, all are related to the covariance matrix (eq.(24)) and are using a 16 point window for averaging. The first estimator is the MDS which is given by the mean over the diagonal elements of eq.(24) when changing *j* and applying a linear regression to obtain *D* and *σ*^2^. The second estimator is the CVE which have *j* = 1 and we solve for the *D, σ*^2^ and take the mean over the diagonal of the covariance matrix. Lastly is the ECVE which is given by eq.(6) in the main text. In Fig.(C.2) the three tests are shown. The CVE has a poor contrast and the two lower diffusion speeds can hardly be distinguished. This is the drawback of using a single step where not enough difference has accumulated. Looking at the MSD, one can identify all three levels but there is a bias for slow diffusion. The ECVE shows better contrast compared to both the CVE and MSD and less bias for the two first diffusion rates. Estimating the positioning error *σ*^2^ for the three methods (see Fig.(C.3)) shows that the MSD has a large bias that is often negatively valued. Both the CVE and ECVE have better performance and have no bias.

#### C. Hidden Markov Model for estimated diffusion traces

Suppose that during a measurement, we obtain a set of measurements of the positions {***r***} and from these we calculate a set of diffusion constants {*D*_*i*_} using the ECVE method described in the main text. Assuming a single diffusion constant, no motion blur, and no localization error, we have a set of diffusion values {*D*_*i*_} which originates as the variance of a normal distribution with a known mean (mean is zero) but an unknown variance. The distribution of the diffusion set {*D*_*i*_} can be assumed to be inverse-Gamma distributed. Consider now a measurement sequence of *M* steps, 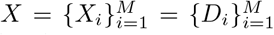, in cells where the underlying dynamics is governed by an HMM model on diffusion rates. The distribution of our measured sequence, 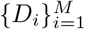, is thus a mixture of inverse-Gamma distributions, with parameters *ϕ* = {*ϕ*_*j*_}, corresponding to different states *S* = {*S*_*j*_} where *j* ∈ {1, …, *N* } iterates over the *N* different states. The relation between the states is given by an *N* × *N* transition matrix ***a*** with elements *a*_*kl*_ which are the probability of moving from state *k* to state *l* with the normalization condition Σ_*l*_ *a*_*kl*_ = 1. Using these we can construct the likelihood of obtaining a sequence of measurements for the unknown parameters *γ* = {***a***, *S, ϕ*}. Now, the goal of the game is to try to find the parameters that maximize the likelihood of the measured sequence. Note that if we did know the state *S*_*j*_ for each measurement *X*_*i*_ we would be able to organize our data in pairs 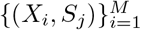 which indicates exactly which data point belong to which state and thus which distribution. The parameter *ϕ*_*j*_ for each distribution can be calculated separately and thus the probabilities *P* (*X*_*i*_|*S*_*j*_) can be obtained from each inverseGamma distribution. The likelihood of obtaining a sequence 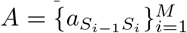 from an unknown transition matrix ***a*** is then given by

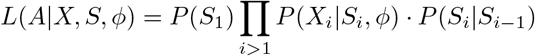

where 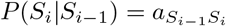, with *P*(*S*_1_) as the probability of starting at state *S*_1_. The problem is thus reduced to maximizing a multidimensional first degree polynomial. Unfortunately the state sequence is not known and are thus considered hidden, to account for this we marginalize over the state *S*, and the likelihood becomes

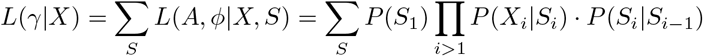

finding 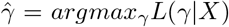 is a much harder problem, where numerical search is required. For this task there are several packages available, here we use the python Pomegranate HMM suite [40].

**Fig. C.1.**
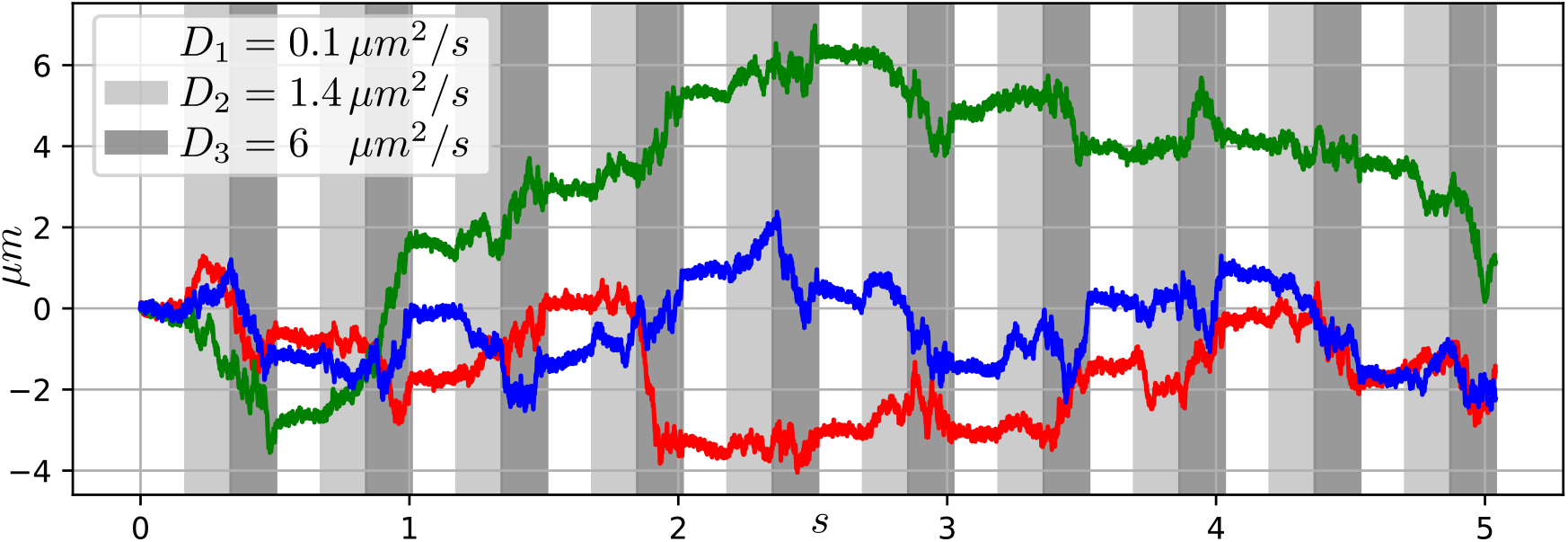
Trajectory coordinates for x (red), y (green) and z (blue). Each vertical section corresponds to 168*ms* and within each section a fixed diffusion rate is maintained. The diffusion rates used are 0.1*µm*^2^*/s* (white), 1.4*µm*^2^*/s* (light gray) and 6*µm*^2^*/s* (dark gray)

**Fig. C.2.**
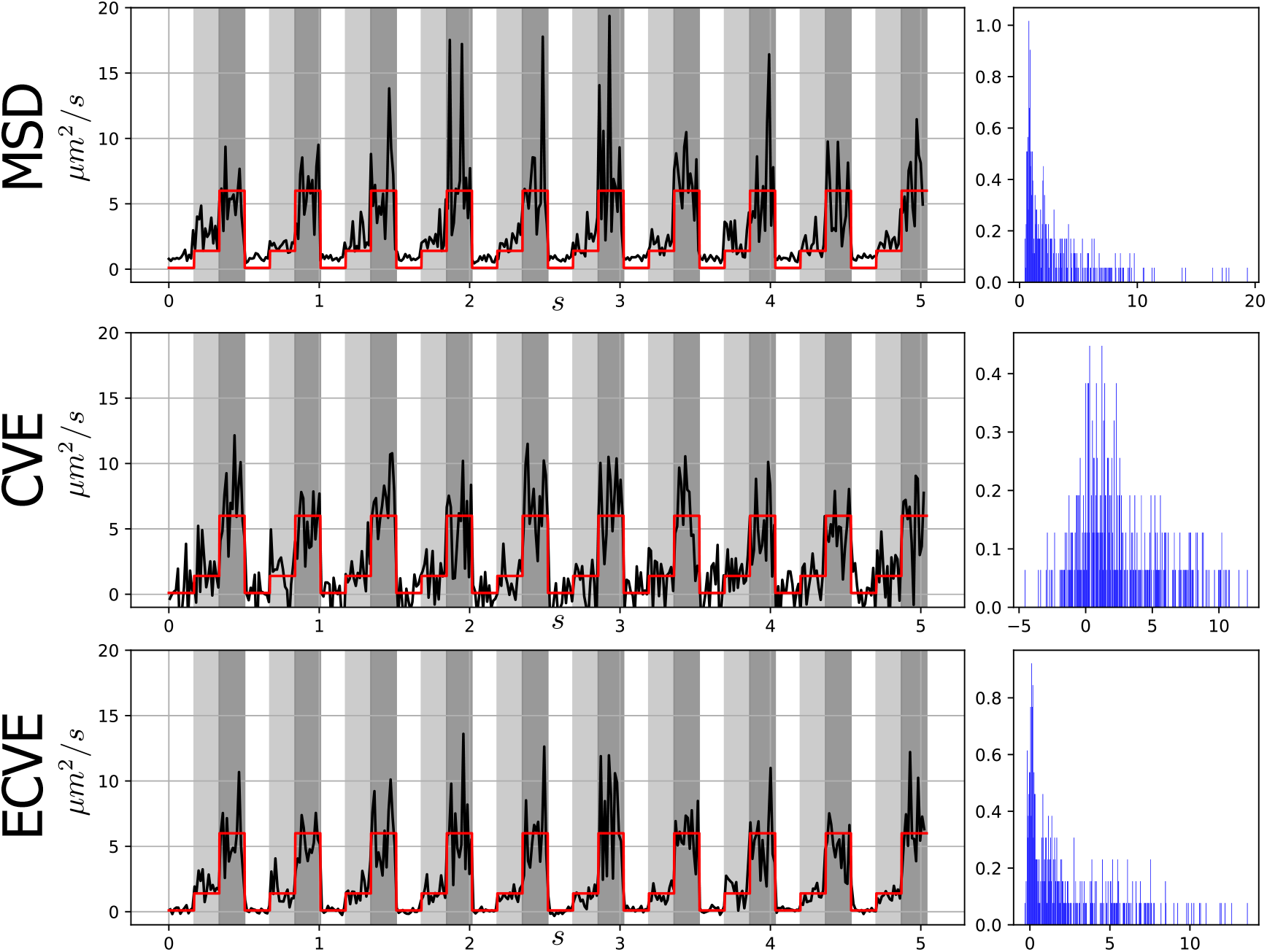
Estimation of the diffusion over the trajectory in Fig.(C.1), red is the ground truth and to the right the histogram of the trajectory.

**Fig. C.3.**
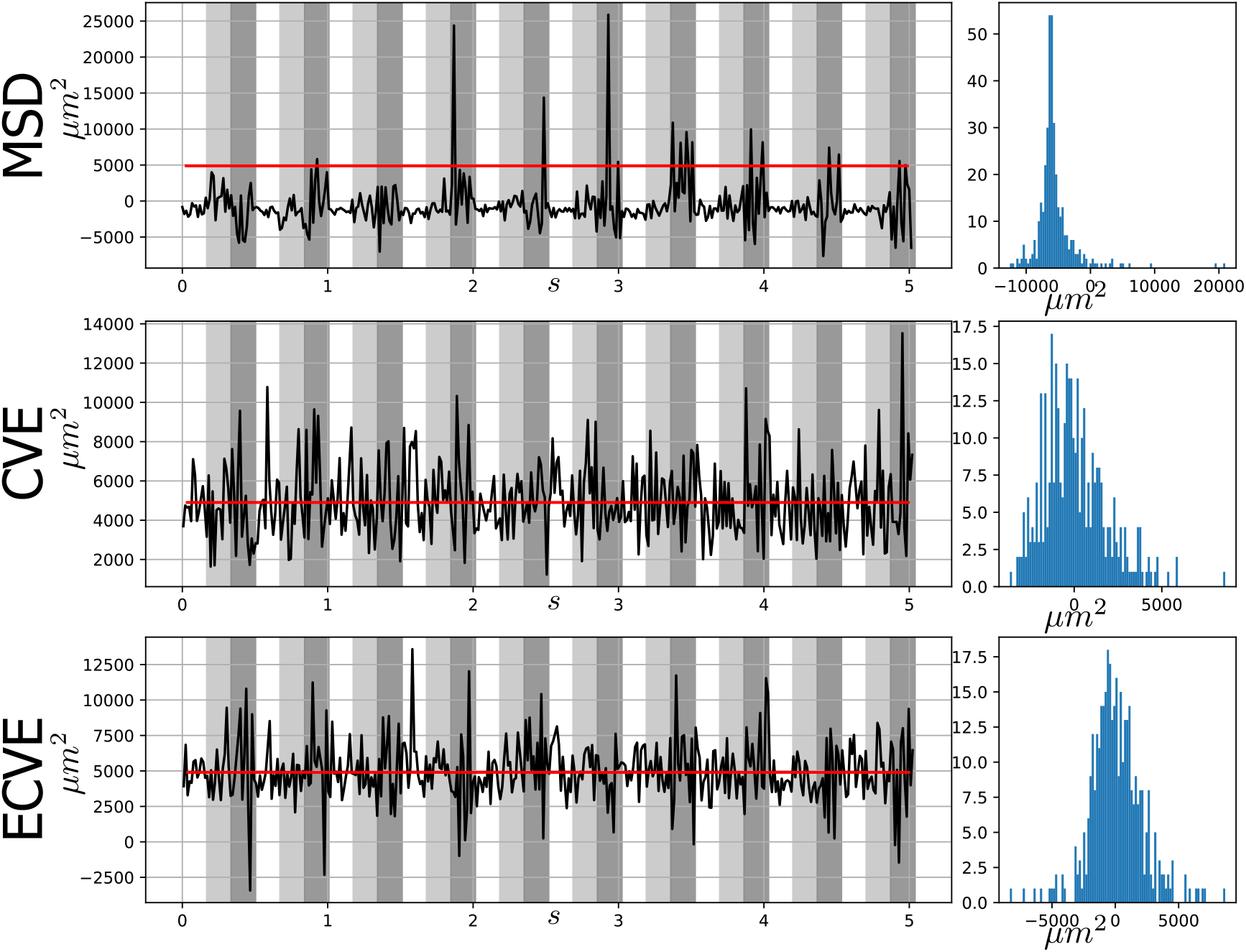
Estimation of *σ*^2^ the trajectory in Fig.(C.1), red is the ground truth and corresponds to a standard deviation of 70*µm*. To the right the histogram of the distance to the ground truth.

##### 1) Gamma and inverse-Gamma distribution relation

Within the Pomegranate HMM suite, the inverse-Gamma distribution is not available but the Gamma distribution is. Fortunately, the two are closely related. The gamma distribution is given by

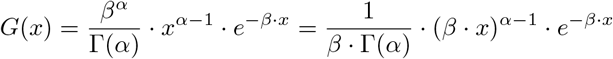

where 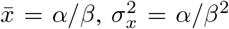We can relate the Gamma distribution to the inverse-Gamma by the transformation *y* = 1*/x*, from probability theory we have that *G*_*y*_(*y*) = *G*_*x*_(1*/y*) · *J* (*x* = 1*/y*), where *J* is the Jacobian of the transformation. This gives

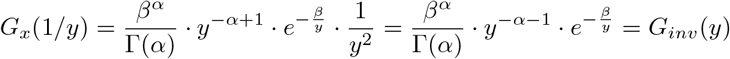

Thus, getting the parameters for the inverse-Gamma distribution is the same as the ones for the Gamma with inverted data. Finally, there is a slightly different interpretation of the parameters: *β* is the rate parameter in the Gamma distribution but is the scale parameter in the inverse-Gamma distribution.

#### D. Handling negative values when fitting inverse-gamma distributions

For all three estimators tested, MSD, ECV and ECVE, one might get negative values. These occur when evaluating diffusion close to zero and will be affected by the noise level in the signal. The negative values will be less when using higher offset *j* compare to |*k* − *k*′|, but this also depends on the dynamics analyzed. One way to handle negative values is to move to higher offset values, *j*, or average over a larger interval. However, this is not desirable if the goal is analysis of fast dynamics.

For a Gaussian distribution with known mean and unknown variance, the most apt distribution over the unknown variance is inverse-Gamma. Unfortunately, the inverse-Gamma is a distribution for positive values, and it would be preferable to be able to handle small negative values within the inverse-Gamma model. The way we handle this is to shift all values by a bias to the positive axis. We note that if we shift the data by the amount *b* such that the data is *x* + *b* we have that 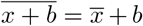 and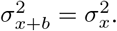.

##### 1) Shifting the inverse-Gamma distribution

The inverse gamma distribution is given by

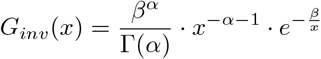

whe*re* 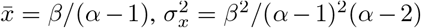if *α >* 2. Thus this means that if we get parameters *α*′ and *β*′ for the {*x* + *b*}data we get 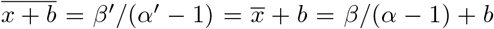 and that 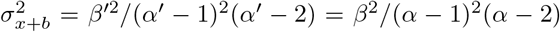 where *α* and *β* are the non shifted parameters. This gives the relations between shifted and non shifted data:

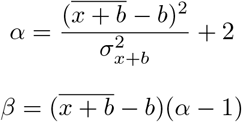

We note that by shifting the inverse-Gamma a positive amount, we are forcing the distribution to be more Gaussian like. One is thus losing the skewness of the inverse-Gamma distribution. The skewness is an important property that we desire to preserve. Thus, the shift must be relatively small. On the other hand, if no shift is added we allow for distributions with infinite variance. By adding a shift, the variance is suppressed. A shift of 0.01 − 10*µm*^2^*/s* gives good results and for most data we used a shift of 2*µm*^2^*/s*. The exception is for *E. coli* Trigger Factor data measured with the pattern which has a missing center shot and is presented in sub.sec.(F), for this data we used a shift of 5*µm*^2^*/s*.

#### E. Bootstrap sampling for error estimation

To estimate error in the HMM parameters for simulations and TF data, we did a bootstrap sampling over the trajectories. From the pool of trajectories we draw with replacement the same number of trajectories as there are in the pool. The new trajectory set is analyzed with the HMM analysis. This is done 100 times, giving us a spread in the HMM parameter estimation.

## Appendix D

### Diffusion model in cell like structure

Simulations are performed by setting up a diffusion HMM model with 3 states characterized by the transition matrix shown in Fig.(D.1). From this, we generate Viterbi paths where the length is selected at random within the interval 50*ms* and 800*ms*. The Viterbi paths are then used for creating particle trajectories where the x,y,z step sizes are sampled from a gaussian with the proper diffusion rate. At this stage we have an unbounded diffusion. By folding this trajectory within a cell structure using reflective boundary conditions, we obtain a diffusion ground truth path within a cell with a time resolution of 20*µs*. For diffusion analysis of simulated trajectories, we used the ECVE with a time lag of *j* = 8 with a 16 point localization time window, see Fig.(D.2**b**). In the main text we present the results when simulating 100 trajectories using the model in Fig.(D.1**a**). We see that the fitted HMM has a hard time extracting the correct parameters for the diffusion rates and occupancies. This also holds for the dwell times which are poorly estimated.

In Fig.(D.2) we present results of the second simulation based on the 3-state HMM in Fig.(D.1**b**). For this model 50 trajectories are simulated in an *E. coli* like geometry. The diffusion estimations from the reconstructed trajectories are fitted to a 3-state HMM assuming inverse-gamma distributions as emission distributions. In Fig.(D.2**b**), the red trace is the predicted Viterbi path, and in Fig.(D.2**c**), we show the distribution of all estimated diffusion values together with the predicted inversegamma distributions from the trained HMM. As before we see that the histogram does not correctly reflect the fast state. Instead, we see a mean value between the fast and the intermediate state. Two major distributions can be identified in the histogram: the slow with mean diffusion 0.26 ± 0.03*µm*^2^*/s* (ground truth 0.1*µm*^2^*/s*), with s.e. from bootstrapping, and occupancy 33 ± 8% (ground truth 30%) and the fast state with mean diffusion 3.18 ± 0.26*µm*^2^*/s* (ground truth 6*µm*^2^*/s*) and occupancy 32 ± 8% (ground truth 37%). The intermediate state is identified by the HMM analysis in between the two distributions with a 36 ± 8% (ground truth 33%) occupancy and mean diffusion 1.45 ± 0.24*µm*^2^*/s* (ground truth 1.4*µm*^2^*/s*). Here, the occupancies are here better estimated, and the dwell times are also better represented with 373*ms* (300*ms*) for the bound state, 44*ms* (30*ms*) for the intermediate state and 47*ms* (30*ms*) for the fast state, where the ground truth is reported within parentheses. The estimated parameters are better but the predicted Viterbi path is still not capturing the faster transitions of a trajectory.

**Fig. D.1.**
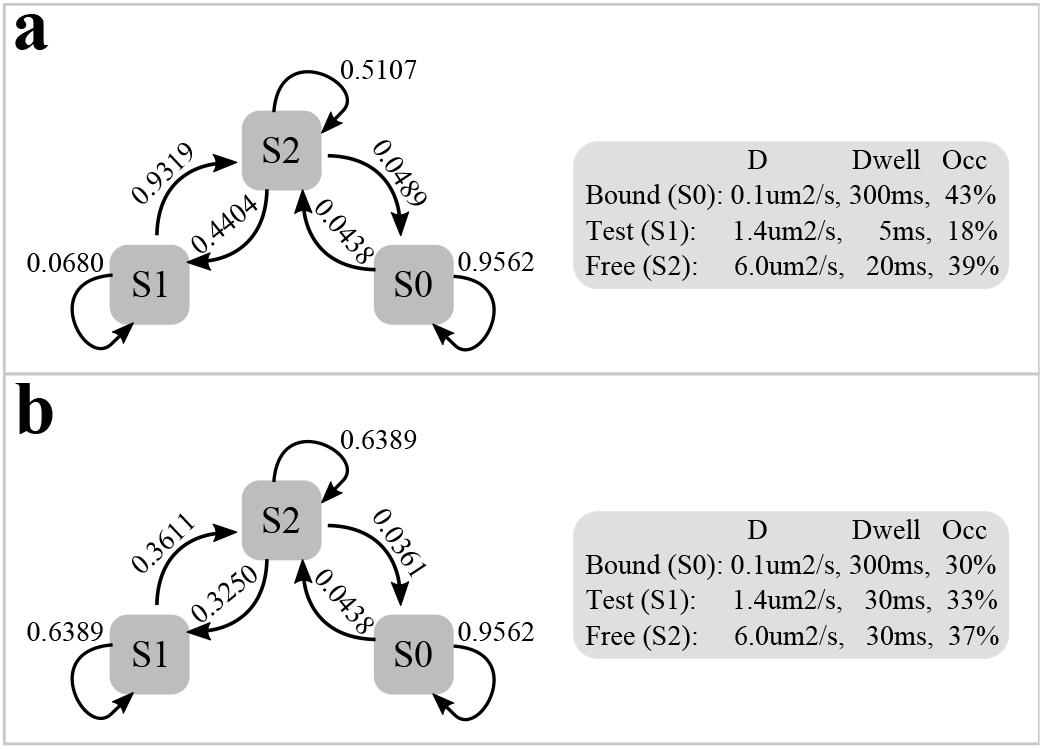
HMM model parameters for simulated trajectories in *E. coli* like geometry, transition probabilities are for a 13.44ms which corresponds to the time window of 16 localizations. **a** is the model with a short-lived intermediate state. **b** is a more relaxed model where the free and intermediate state have both a dwell time of 30ms.

**Fig. D.2.**
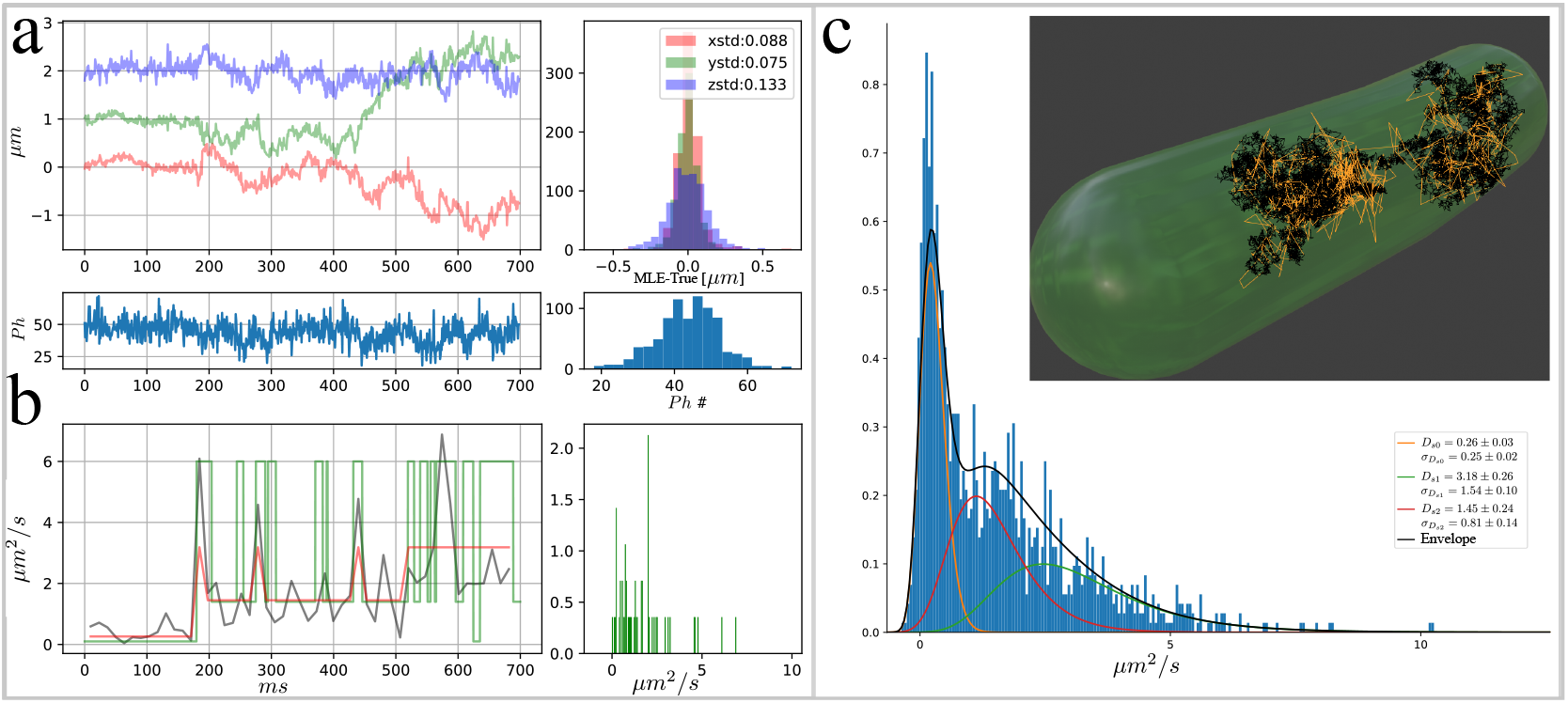
Example trajectory with HMM analysis of all 50 simulated trajectories of the model in Fig.(D.1**b**). **a**, upper left; example trajectory with x (red) y (green) z (blue) estimated position coordinates with corresponding photon counts (bottom). Upper right; the deviation away from ground truth with the standard deviation of the error distribution in the legend, (bottom) is the photon count distribution. **b**, left; diffusion point estimation (gray) along the trajectory in **a**, ground truth diffusion is in green, and the HMM prediction in red. Right; the distribution of estimated diffusions over the trajectory in **a. c**, distribution of all diffusion estimates from the 50 trajectories. The 3 state HMM model prediction of the emission distributions are plotted on top of the histogram. The inverse-gamma mean and standard deviation with s.e. of each emission distribution is reported in the legend. Inset, 3D representation of the cell volume with ground truth trajectory in black and the estimated path from **a** in yellow.

## Appendix E

### Tracking validation

Using an experimentally measured PSF z-stack that has been interpolated down to voxels of 10nm, we simulate the microscope laser pattern with an added gaussian position noise of 10nm for xy and 10nm for z. The gaussian noise is added since characterization of the beam profile after reflection on a reflective sample indicates a small beam/sample movement at high frame rates (200fps) when analyzing images taken from a EMCCD. At low frame rate, this is averaged out; however, for tracking at 1ms time scale, this must be taken into account.

From the PSF stack we sample the PSF intensities and generate photon counts assuming Poissonian statistics. The real-time feedback loop for the piezo stage is simulated as a long moving average over the ground truth data. This limits the speed of the simulated piezo-stage to behave close to what is observed in experiments. The output of the simulation is of the same format as a real experiment and thus can be analyzed by the same pipeline as experimental acquired data.

#### A. Rotational motion

Validation of the tracking is done by comparing simulation and real-time tracking of beads. For these we simulated similar rotational motion and at photon counts similar to experiments. A comparison between tracking of beads and simulations is shown in main Fig.(2) and a measurement at higher photon rate is shown in Fig.(E.1).

**Fig. E.1.**
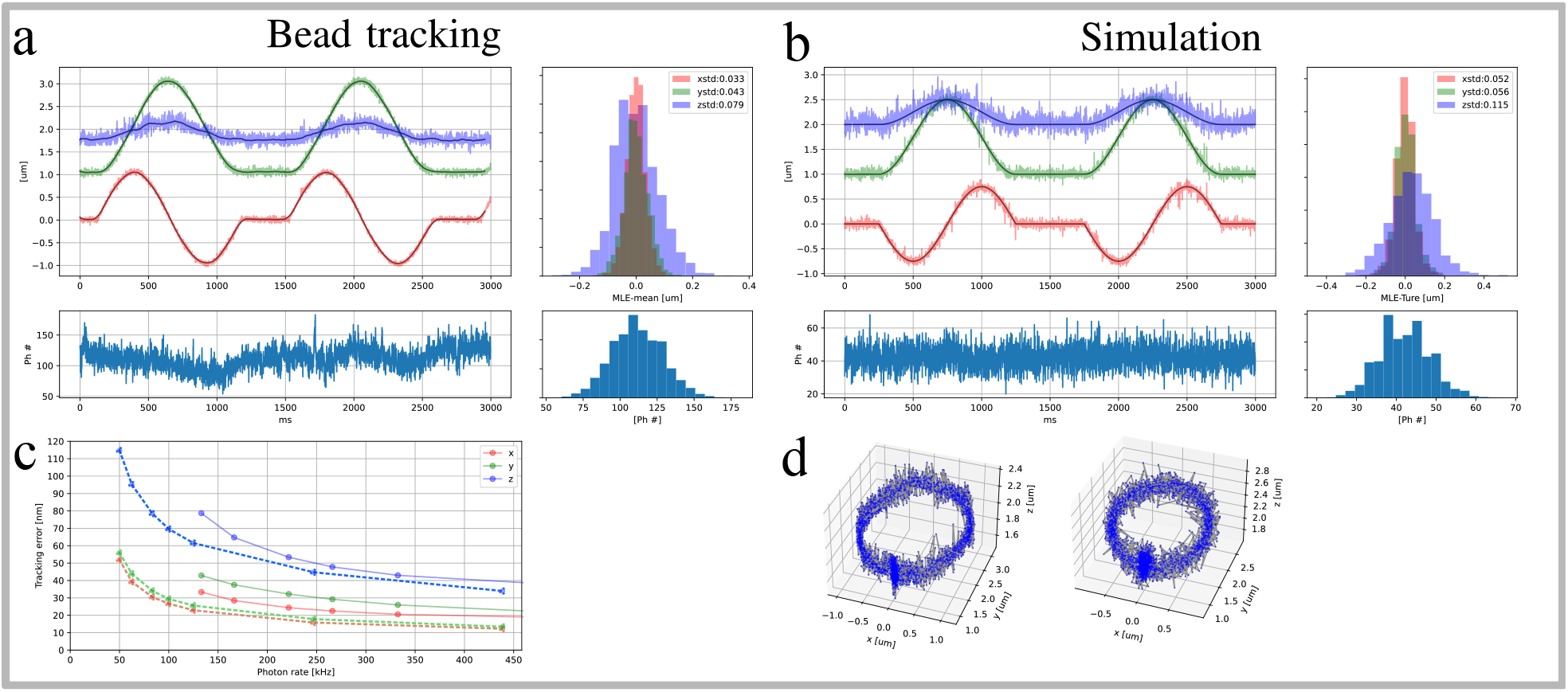
Resolution test at a higher photon count rate. **a**, Tracking of beads fixed in agarose, the piezo stage is programmed to trace circles with a x,y-radius of 1*µm* and a z peak to peak oscillation of 0.5*µm*. Upper left; Trajectory estimated x (red), y (green) and z (blue) coordinates with a temporal resolution of 0.84*ms*. Upper right; distribution of standard deviations over the trajectory between estimation and a moving average with a window size of 100 points. Bottom left; photon counts per localization which corresponds to 60kHz photon count rate. To the right the photon count distribution. **b**, Similar to **a** but for a simulated trajectory with an x,y-radius of 0.75*µm* and a z peak to peak oscillation of 0.5*µm*, the error distribution (upper right) is here the distance to ground truth. **c**, resolution as a function of photon count rate for x (red), y (green) and z (blue), solid lines are for the bead data and dashed lines correspond to the simulations. Each point is the standard deviation of the distance histogram in **a** and **b**. These are obtained by increasing the exponential moving average of the likelihood. **d**, 3D point scatter plot of the estimated (left) and simulated (right) trajectory.

To see more clearly the stage behavior, we can take a moving average of the likelihood as described in sec.(C-A4). With the exponential parameter set to 0.4, we get an effective time resolution of 1.98*ms*. In Fig.(E.2) we show the results of the trajectory reconstruction when tracking a bead while the stage is following circle. Upon a closer look at Fig.(E.2**a**), where we have added a 20 point sliding window to give a clearer view, we can identify oscillations and stage corrections. We attribute these artifacts to the stage feedback system and vibrations induced by the movement.

## Appendix F

### Removing the center shot

The theory predicts that higher resolution can be obtained by moving the Gaussian beams further apart. This means that removing the center position in our pattern should increase the tracking resolution when keeping the same photon counts. To test this we did all the measurements with a pattern configuration where the center shot is removed, in practice this means that there is a hole in the center of our pattern. The resolution test in fig(F.1) shows that the simulation loses resolution compared to Fig.(E.1). What can be observed is that the pattern with a hole in the center has more sporadic jumps in the reconstruction indicating that the removal of the center lowers the informational content and thus lowers the noise tolerance. The bead tracking experiment does indicate a small resolution increase in the lateral direction but this is marginal. Looking at higher averaging (Fig.(F.2)) we see, as before, oscillations when tracking while the stage is moving, again we attribute these to the stage feedback-loop and induced vibrations during stage movement.

As for the original pattern, we used the new pattern to simulate on an *E. coli*-like geometry and experimental measurements on the *E. coli* Trigger Factor. Results of the simulation when generating 100 trajectories according to the HMM model apper in Fig.(D.1**a**) and use the same analysis as for the data reported in the main text. The diffusion estimations from the reconstructed trajectories are fitted to a 3-state HMM assuming inverse-gamma distributions as emission distributions. In Fig.(F.3**b**), the red trace is the predicted Viterbi path, and in Fig.(F.3**c**), we show the distribution of all estimated diffusion values together with the predicted inverse-gamma distributions from the trained HMM. As before we see that the histogram does not correctly reflect the fast state, we instead see a mean value between the fast and the intermediate state. Two major distributions can be identified in the histogram: the slow with mean diffusion 0.21 ± 0.02*µm*^2^*/s* (ground truth 0.1*µm*^2^*/s*), with s.e., and occupancy ± 7% (ground truth 43%) and the fast state with mean diffusion 2.90 ± 0.05*µm*^2^*/s* (ground truth 6*µm*^2^*/s*) and occupancy ± 4% (ground truth 38%). The intermediate state is identified by the HMM analysis in between the two distributions with a 17 ± 5% (ground truth 18%) occupancy and mean diffusion 0.60 ± 0.14*µm*^2^*/s* (ground truth 1.4*µm*^2^*/s*). As seen, the results are similar to those presented with the full pattern. The occupancy is about right, the slow state is clear in the histogram, but we see an average of the fast and intermediate state for the diffusion rate.

**Fig. E.2.**
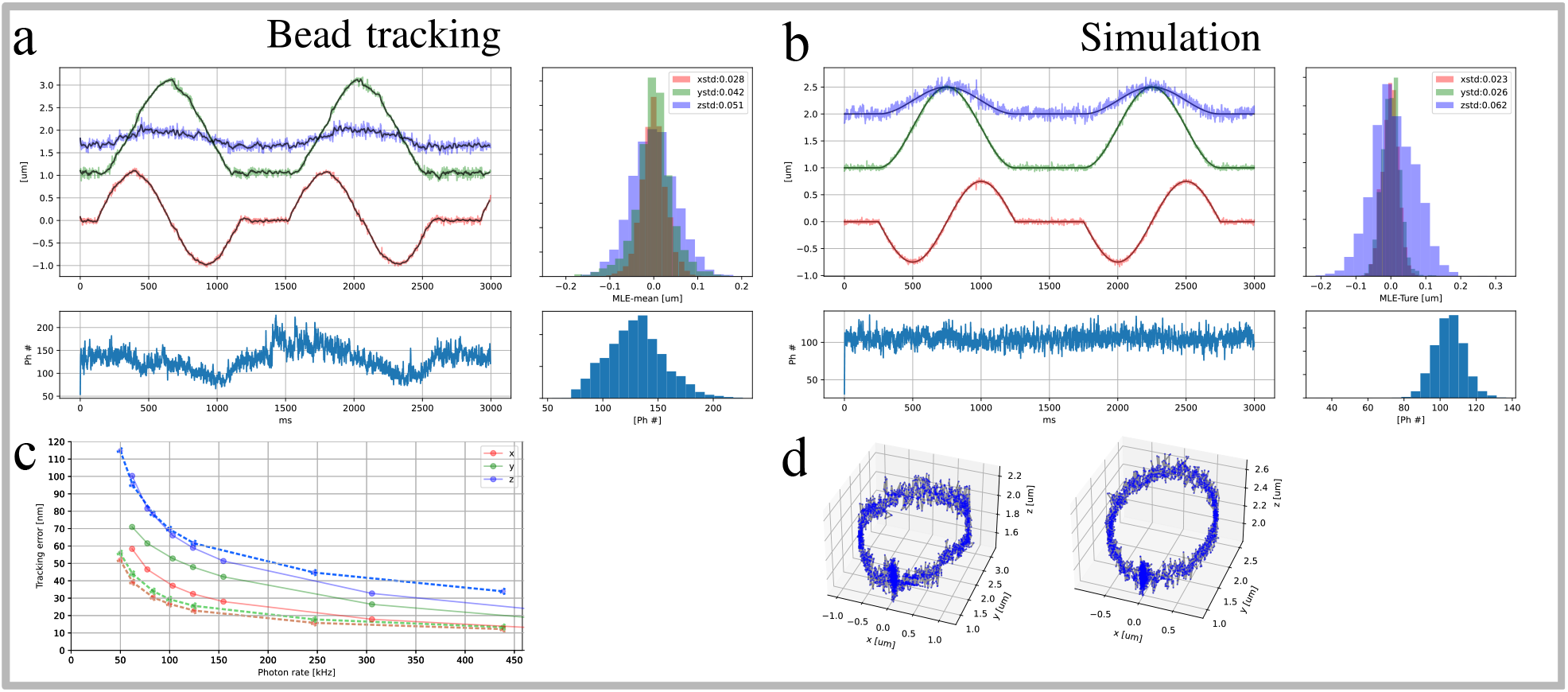
Resolution test when taking a moving average during trajectory reconstruction giving an effective time resolution of 1.98*ms*. **a**, Tracking of beads fixed in agarose, the piezo stage is programmed to follow circles with a x,y-radius of 1*µm* and a z peak to peak oscillation of 0.5*µm*. Upper left; Trajectory estimated x (red), y (green) and z (blue) coordinates with a 0.84*ms* between localizations. Upper right; distribution of standard deviations over the trajectory between estimation and a moving average with a window size of 20 points. Bottom left; photon counts per localization which corresponds to 60kHz photon count rate. To the right the photon count distribution. **b**, Similar to **a** but for a simulated trajectory with an x,y-radius of 0.75*µm* and a z peak to peak oscillation of 0.5*µm*, the error distribution (upper right) is here the distance to ground truth. **c**, resolution as a function of photon count rate for x (red), y (green) and z (blue), solid lines are for the bead data and dashed lines correspond to the simulations. Each point is the standard deviation of the distance histogram in **a** and **b**.These are obtained by increasing the exponential moving average of the likelihood. **d**, 3D point scatter plot of the estimated (left) and simulated (right) trajectory.

**Fig. F.1.**
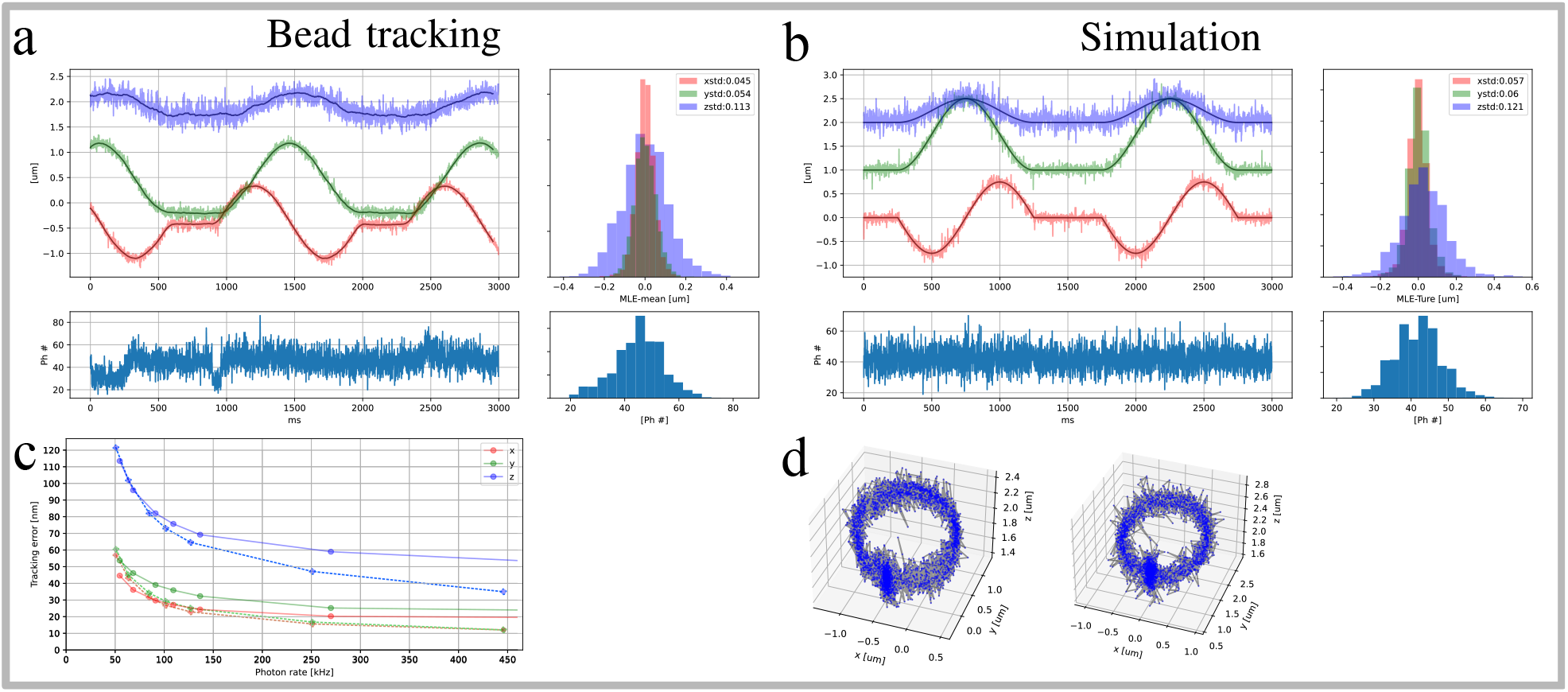
Resolution test when removing the center shot in the tracking pattern. **a**, Tracking of beads fixed in agarose, the piezo stage is programmed to trace circles with a x,y-radius of 1*µm* and a z peak to peak oscillation of 0.5*µm*. Upper left; Trajectory estimated x (red), y (green) and z (blue) coordinates with a 0.82*ms* between localizations (41 shots in this pattern). Upper right; distribution of standard deviations over the trajectory between estimation and a moving average with a window size of 100 points. Bottom left; photon counts per localization which corresponds to 60kHz photon count rate. To the right the photon count distribution. **b**, Similar to **a** but for a simulated trajectory with an x,y-radius of 0.75*µm* and a z peak to peak oscillation of 0.5*µm*, the error distribution (upper right) is here the distance to ground truth. **c**, resolution as a function of photon count rate for x (red), y (green) and z (blue), solid lines are for the bead data and dashed lines correspond to the simulations. Each point is the standard deviation of the distance histogram in **a** and **b**.These are obtained by increasing the exponential moving average of the likelihood. **d**, 3D point scatter plot of the estimated (left) and simulated (right) trajectory.

Measurements on *E. coli* Trigger Factor were also done using the pattern with a hole at the center. In Fig.(F.4) the results are presented; both are selected to show the dynamical behavior of the WT and mutant TF and for that reason, they are longer than the average trajectory. In the case of the wild-type, see Fig.(F.4**a**), we frequently observe a bound state at low diffusion which is interrupted by occasional periods of high diffusion. All trajectories are fitted to a 3-state HMM, see Fig.(F.4**a**) right side where the identified HMM is plotted together with the diffusion histogram. The occupancies for the bound (0.43 ± 0.05*µm*^2^*/s*)and the intermediate (1.20 ± 0.15*µm*^2^*/s*) states are 21 ± 5% and 48 ± 2% respectively and constitute the majority of the events. The parantheses and the HMM errors are from bootstraping over trajectories 100 times. The remaining 30 ± 5% of the time, the TF is in a more free state (2.80 ± 0.14*µm*^2^*/s*) exploring a larger volume. Looking at the mutant, we see a very different situation, see Fig.(F.4**b**). A bound state (1.02 ± 0.09*µm*^2^*/s*) is found by the HMM, with an occupancy of 22 ± 6%. The mutant’s intermediate state (2.36 ± 0.12*µm*^2^*/s*) is here as fast as the wild-type’s free state and is occupied 38 ± 6% of the time. As before we see in the histogram a large tail at higher diffusion which the HMM assigns to a fast (4.60 ± 0.20*µm*^2^*/s*)state with an occupancy of 40 ± 6%. It is clear that the mutant’s mobility is much higher than that of the wild-type, allowing the mutant TF to explore larger parts of the *E. coli* cell. In conclusion the diffusion speeds available for the mutant TF are higher than those allowed for the wild-type TF which was also seen with the tracking pattern with the center shot.

**Fig. F.2.**
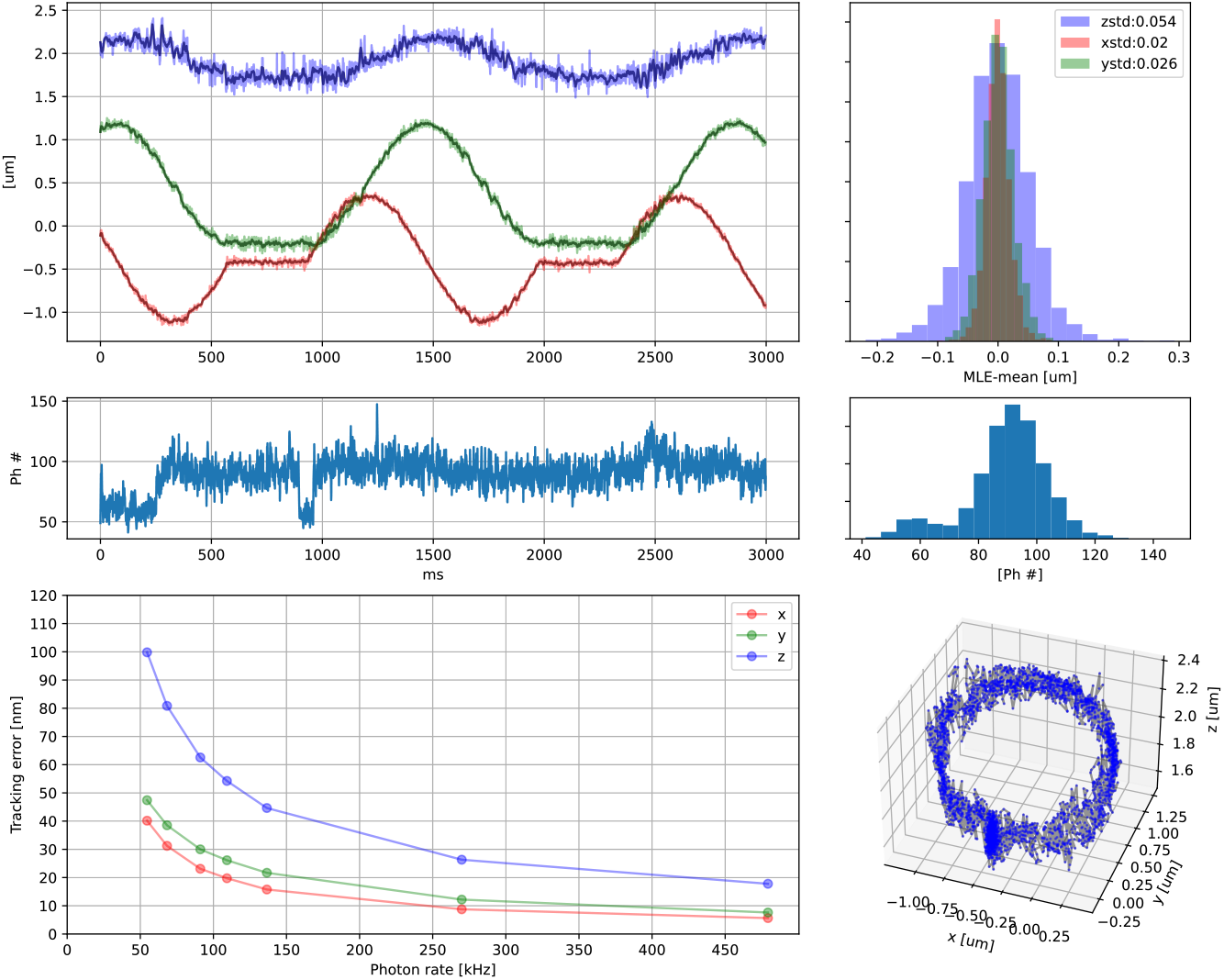
Tracking beads fixed in agarose, the piezo stage was programmed to follow 1Hz circles of x,y-radius of 1*µm* and a z peak to peak oscillation of 0.5*µm*. During reconstruction a moving average is used with the averaging parameter 0.4 giving an effective time resolution of 1.93*ms*. Upper left; Trajectory estimated x (red), y (green) and z (blue) coordinates. Clear oscillations can be seen in the x and y trajectory. Upper right; distribution of standard deviations over the trajectory between estimation and a moving average with a window size of 10 points. Middle left; photon counts per localization and to the right its distribution. Bottom left; resolution as a function of photon rate for x (red), y (green) and z (blue). Obtained by increasing the exponential moving average of the likelihood. Bottom right; 3D point scatter plot of the estimated trajectory.

**Fig. F.3.**
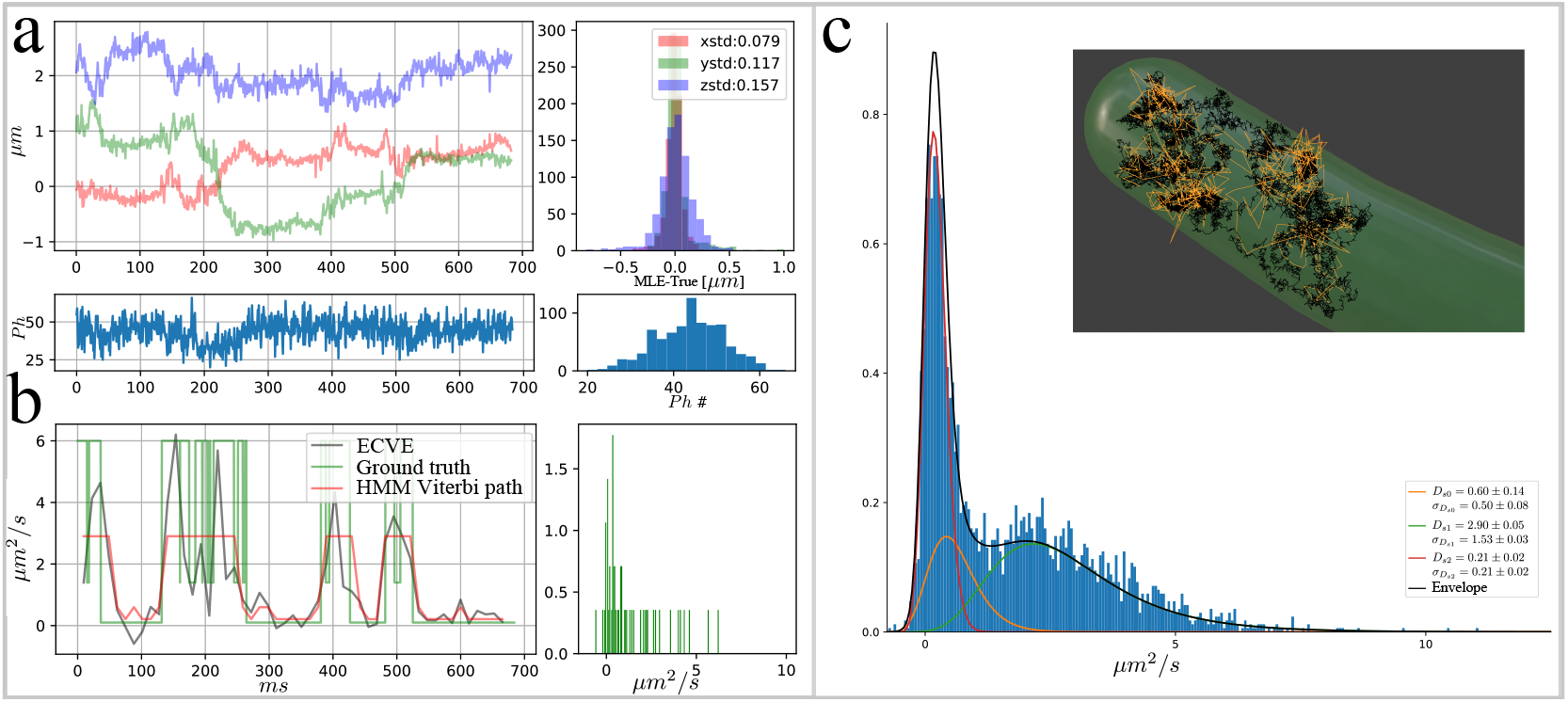
Example trajectory with HMM analysis of all 100 simulated trajectories with a shot pattern with a hole at the center. **a**, upper left; example trajectory with x (red) y (green) z (blue) estimated position coordinates with corresponding photon counts (bottom). Upper right; the deviation away from ground truth with the standard deviation of the error distribution in the legend, (bottom) is the photon count distribution. **b**, left; diffusion point estimation (gray) along the trajectory in **a**, ground truth diffusion is in green, and the HMM prediction in red. Right; the distribution of estimated diffusions over the trajectory in **a. c**, distribution of all diffusion estimates from 100 trajectories. The 3 state HMM model prediction of the emission distributions are plotted on top of the histogram, the inverse-gamma mean and standard deviation with s.e. of each emission distribution is reported in the legend. Inset, 3D representation of the cell volume with ground truth trajectory in black and the estimated path from **a** in yellow.

**Fig. F.4.**
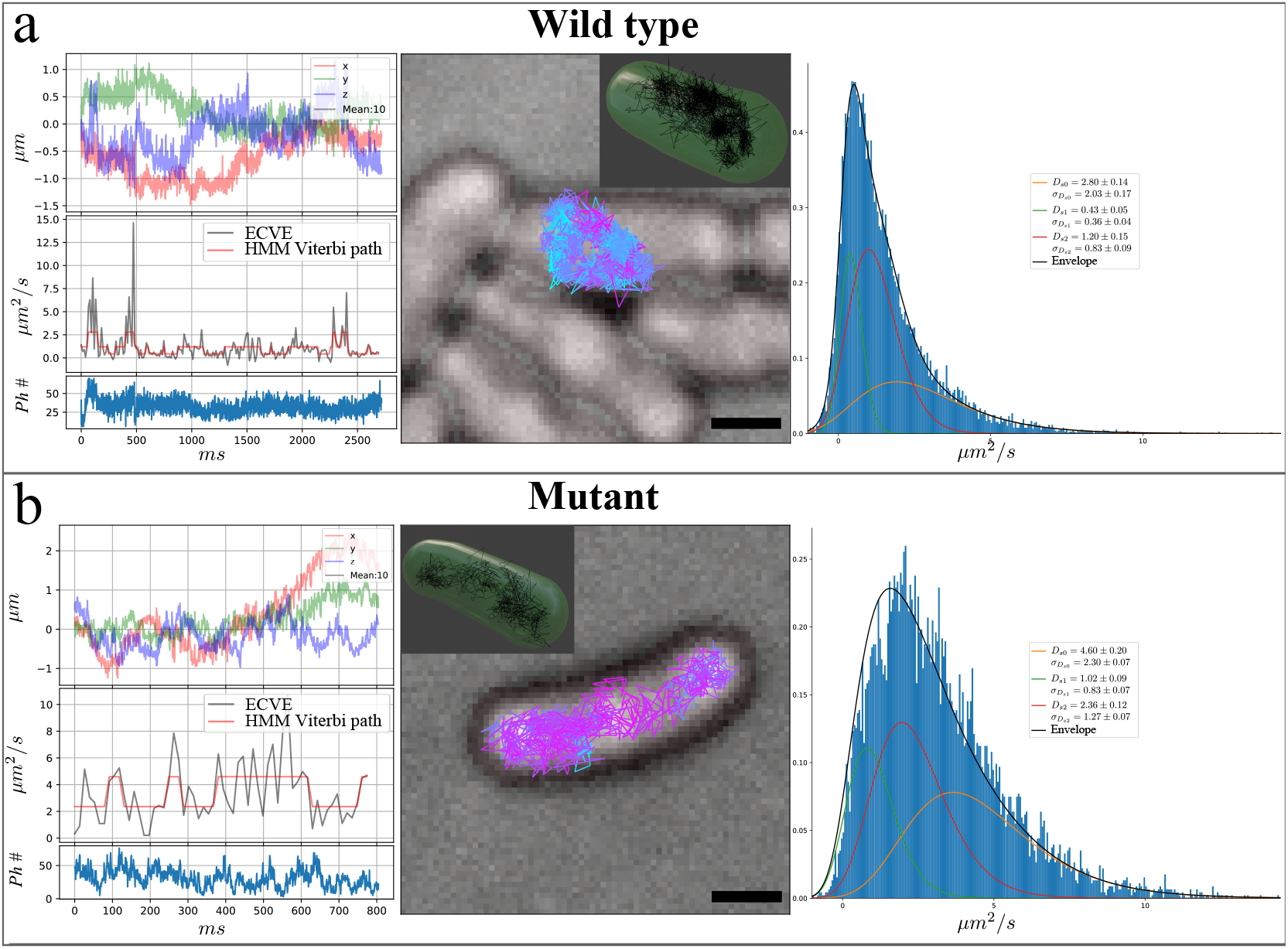
For each sample type, wild-type **a** and mutant **b**, an example trajectory (left) is shown together with the distribution of the accumulated diffusion histograms over all measured trajectories of each type (right). Each horizontal block shows: center image, widefield cell image (black bar 1µ*m*) with estimated trajectory color coded on diffusion estimation. Inset is the 3D trajectory with a cell membrane representation. Left plots, for the same trajectory shown in the center, are from the top; position estimation, point wise diffusion estimation (gray) with HMM Viterbi path (red) and photon counts in the bottom plot. Right plot; histograms over all diffusion estimates acquired in each measurement together with the HMM emission distributions. In the histogram legend are the found mean diffusion values and distribution standard deviation with s.e esstimated by bootstraping 100 iterations over the trajectories. For the wild-type 319 trajectories were acquired from 85 cells. And for the mutant 565 trajectories were acquired from 53 cells.

## Appendix G

### Trajectory length distributions for Trigger Factor experiments

The trajectory length distributions are found in Fig.(G.1). To avoid losing tracking events we run the real time tracking with a release threshold that occasionally is tracking background. In the post-processing, we apply an HMM on the photon counts with a running mean to identify bleaching steps. These bleaching steps are used to truncate trajectories and isolate single molecule tracking events.

## Appendix H

### Sample preparations

#### A. Bead sample

Agarose was dissolved in DI to make 4% gel. After heating the gel to 80^*°*^*C*, 20*nm* TetraSpeck beads were added to a final concentration of 500*pM*. The bead-gel mixture was poured to form a pad between a clean glass slide and quartz coverslip.

#### B. Strain construction and sample preparation

Tracking of TF-HaloTag was performed in *Escherichia coli MG*1655Δ*tig* in which *tig*, encoding Trigger Factor, was deleted from the chromosome by λ Red assisted recombineering [41] and replaced by a kanamycin resistance marker. The strain carries either the pQE30lacIq_tig_halotag plasmid in the case of tracking WT TF-HaloTag, or pQE30lacIq_tig_halotag_FRK_AAA_mut in the case of tracking the TF-HaloTag mutant. Without IPTG induction, the plasmids show leaky expression of TF-HaloTag (data not shown).

**Fig. G.1.**
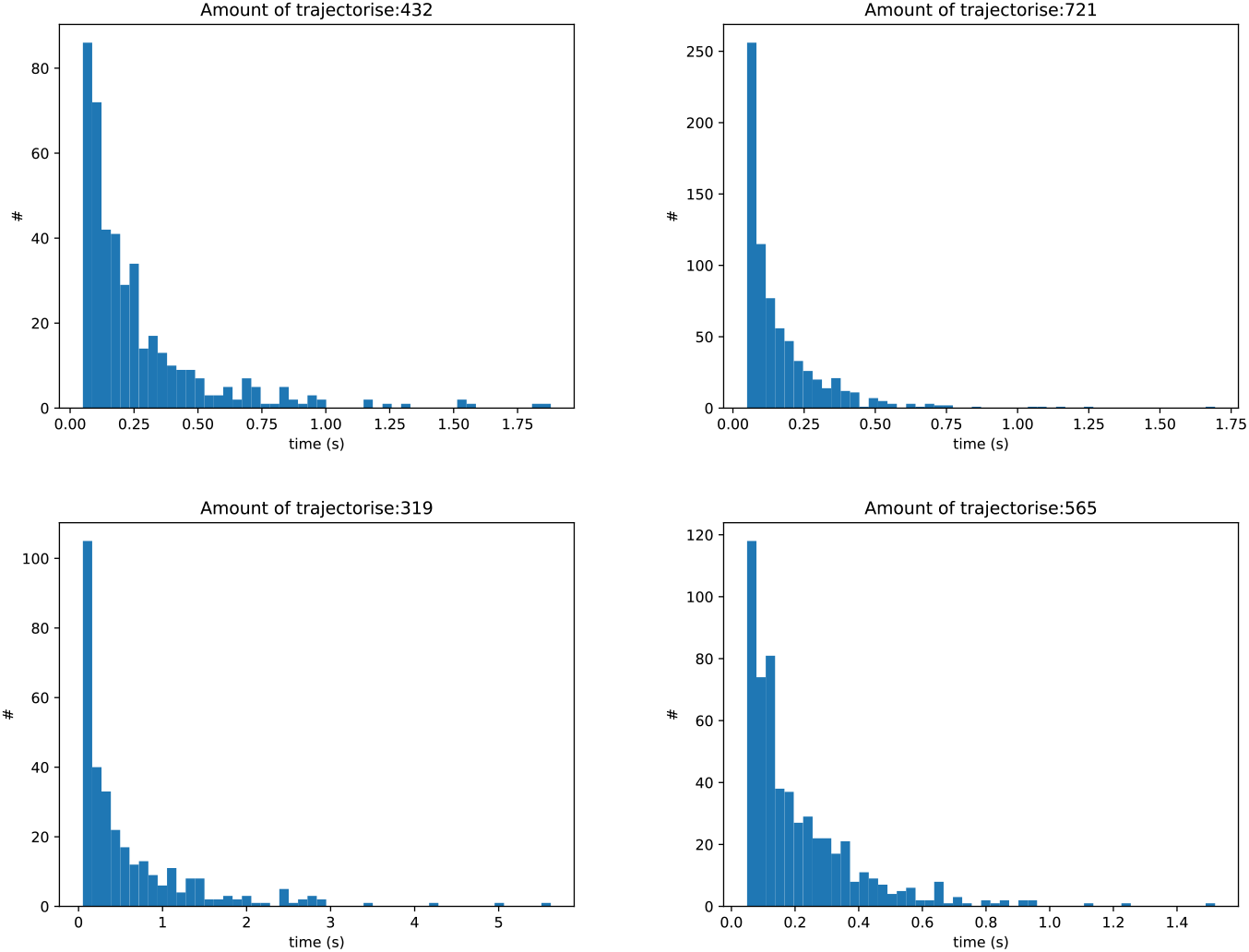
Trajectory length distributions, upper row shows the experiment presented in the main article, bottom row is for the experiment when removing the center shot which is presented in sub. sec. (F). Left, wild-type with the total amount of trajectories in the title. Right side are the distributions for the mutant.

Cells from a glycerol stock were inoculated in Luria Broth (LB), 50 *µg/ml* kanamycin and 100 *µg/ml* carbenicillin and incubated at 37°*C* 200 rpm overnight. The overnight culture was diluted 1:100 in 10 ml fresh LB supplemented with antibiotics and grown at 37°*C* with shaking until *OD*_600_ reached ca 0.5. Cells were harvested at 4000 g, washed in 1 ml M9 media supplemented with 0.2% glucose and resuspended in 150 *µl* EZ Rich Defined Medium (RDM, Teknova) supplemented with 0.2% glucose and 0.1 *µM* Janelia Fluor-549 HaloTag ligand. Cells were labeled for 30 min at 25^*°*^*C*. Excess dye was washed off by adding 1 ml M9 followed by pelleting and resuspension in 1 ml M9. The washing was repeated twice, followed by incubation in 2 ml RDM at 37^*°*^*C* with shaking for 60 min. The washing step was repeated three times and finally cells were resuspended in RDM to an *OD*_600_ of 0.03. The cell suspension was sparsely spread onto a 2% agarose (SeaPlaque GTG Agarose, Lonza) pad in RDM which had been prepared on a 76×32 mm microscopy slide (VWR) with a 1.7×32.8 cm Gene Frame (ThermoFisher Scientific) attached and covered with a 24×32 mm high precision cover glass (ThorLabs).

#### C. Sample measurement routine

All data acquired from cell samples are done by first calibrating the microscope by optimizing the PSF and measuring the PSF. The bead sample is then mounted and tracking of beads is done to see that the microscope is performing as expected. After the calibration the cell sample is mounted, moving through the sample colonies coordinates are registered and a widefield image is taken. Each colony is then re-visited after some time to verify that the cells are growing. At this stage a new widefield image is acquired together with a fluorescent image to verify that fluorescence is present. The sample is scanned before turning on the real-time tracking. All data is saved before moving to the next cell colony. Each experiment constitute of measuring both the WT and the mutant with the same microscope calibration one after the other.

## References

1. English, B. P. et al. Single-molecule investigations of the stringent response machinery in living bacterial cells. Proceedings of the National Academy of Sciences of the United States of America 108, 365–373. ISSN: 00278424 (2011).

2. Elf, J. & Barkefors, I. Single-Molecule Kinetics in Living Cells. Annual Review of Biochemistry 88, 635–659. ISSN: 15454509. 1809.03265 (2019).

3. Gu, L. et al. Molecular resolution imaging by repetitive optical selective exposure. Nature Methods 16, 1114–1118. ISSN: 1548-7105. http://dx.doi.org/10.1038/s41592-019-0544-2 (2019).

4. Reymond, L. et al. SIMPLE: Structured illumination based point localization estimator with enhanced precision. Optics Express 27, 24578. ISSN: 1094-4087 (2019).

5. Cnossen, J. et al. Localization microscopy at doubled precision with patterned illumination. Nature Methods17. ISSN: 1548-7105. http://dx.doi.org/10.1038/s41592-019-0657-7 (2020).

6. Jouchet, P. et al. molecules with modulated excitation. Nature Photonics15. ISSN: 1749-4893. http://dx.doi.org/10.1038/s41566-020-00749-9 (2021).

7. Balzarotti, F. et al. Nanometer resolution imaging and tracking of fluorescent molecules with minimal photon fluxes. Science 612, 606–612. ISSN: 0036-8075. 1611.03401. http://arxiv.org/abs/1611.03401 (2017).

8. Eilers, Y., Ta, H., Gwosch, K. C., Balzarotti, F. & Hell, S. W. MINFLUX monitors rapid molecular jumps with superior spatiotemporal resolution. Proceedings of the National Academy of Sciences of the United States of America 115, 6117–6122. ISSN: 10916490 (2018).

9. Masullo, L. A. et al. Pulsed interleaved MINFLUX. Nano Letters 21, 840–846 (2021).

10. Levi, V., Ruan, Q. Q. & Gratton, E. 3-D particle tracking in a two-photon microscope: Application to the study of molecular dynamics in cells. Biophysical Journal 88, 2919–2928. ISSN: 00063495. http://dx.doi.org/10.1529/biophysj.104.044230 (2005).

11. Berglund, A. J. & Mabuchi, H. Tracking-FCS: Fluorescence correlation spectroscopy of individual particles. Optics Express 13, 8069. ISSN: 1094-4087 (2005).

12. Enderlein, J. Tracking of fluorescent molecules diffusing within membranes. Applied Physics B: Lasers and Optics 71, 773–777. ISSN: 09462171 (2000).

13. Marklund, E. et al. DNA surface exploration and operator bypassing during target search. Nature 583, 858–861. ISSN: 14764687. http://dx.doi.org/10.1038/s41586-020-2413-7 (2020).

14. Hou, S., Lang, X. & Welsher, K. Robust real-time 3D single-particle tracking using a dynamically moving laser spot. Optics Letters 42, 2390. ISSN: 0146-9592 (2017).

15. Hou, S. & Welsher, K. An Adaptive Real-Time 3D Single Particle Tracking Method for Monitoring Viral First Contacts. Small 15, 1–9. ISSN: 16136829 (2019).

16. Perillo, E. P. et al. Deep and high-resolution three-dimensional tracking of single particles using nonlinear and multiplexed illumination. Nature Communications6. ISSN: 20411723 (2015).

17. Hell, S. W. & Wichmann, J. Breaking the diffraction resolution limit by stimulated emission: stimulated-emission-depletion fluorescence microscopy. Optics Letters 19, 780. ISSN: 0146-9592 (1994).

18. Klar, T. A., Jakobs, S., Dyba, M., Egner, A. & Hell, S. W. Fluorescence microscopy with diffraction resolution barrier broken by stimulated emission. Proceedings of the National Academy of Sciences of the United States of America 97, 8206–8210. ISSN: 00278424 (2000).

19. Rust, M. J., Bates, M. & Zhuang, X. Sub-diffraction-limit imaging by stochastic optical reconstruction microscopy (STORM). Nature Methods 3, 793–795. ISSN: 15487091 (2006).

20. Betzig, E. et al. Imaging intracellular fluorescent proteins at nanometer resolution. Science 313, 1642–1645. ISSN: 00368075 (2006).

21. Levi, V., Ruan, Q., Kis-Petikova, K. & Gratton, E. Scanning FCS, a novel method for three-dimensional particle tracking. Biochemical Society Transactions 31, 997–1000. ISSN: 03005127 (2003).

22. Gwosch, K. C. et al. MINFLUX nanoscopy delivers 3D multicolor nanometer resolution in cells. Nature Methods 17, 217–224. ISSN: 15487105. http://dx.doi.org/10.1038/s41592-019-0688-0 (2020).

23. Schmidt, R. et al. MINFLUX nanometer-scale 3D imaging and microsecond-range tracking on a common fluorescence microscope. Nature Communications 12, 1478. ISSN: 2041-1723. https://doi.org/10.1038/s41467-021-21652-z (2021).

24. Berglund, A. J. Statistics of camera-based single-particle tracking. Physical Review E - Statistical, Nonlinear, and Soft Matter Physics 82, 1–8. ISSN: 15393755 (2010).

25. Michalet, X. & Berglund, A. J. Optimal diffusion coefficient estimation in single-particle tracking. Physical Review E - Statistical, Nonlinear, and Soft Matter Physics 85, 1–14. ISSN: 15393755 (2012).

26. Shuang, B. et al. Improved analysis for determining diffusion coefficients from short, single-molecule trajectories with photoblinking. Langmuir 29, 228–234. ISSN: 07437463 (2013).

27. Liu, K., Maciuba, K. & Kaiser, C. M. The Ribosome Cooperates with a Chaperone to Guide Multi-domain Protein Folding. Molecular Cell 74, 310–319.e7. ISSN: 10974164. https://doi.org/10.1016/j.molcel.2019.01.043 (2019).

28. Yang, F. et al. Single-molecule dynamics of the molecular chaperone trigger factor in living cells. Molecular Microbiology 102, 992–1003. ISSN: 13652958. https://www.ncbi.nlm.nih.gov/pmc/articles/PMC5161668/ (2016).

29. Grimm, J. B., Brown, T. A., English, B. P., Lionnet, T. & Lavis, L. D. Synthesis of Janelia Fluor HaloTag and SNAP-Tag Ligands and Their Use in Cellular Imaging Experiments. Methods in Molecular Biology (Clifton, N.J.) 1663, 179–188 (2017).

30. Los, Georgyi V. et al. HaloTag: A Novel Protein Labeling Technology for Cell Imaging and Protein Analysis. American Journal of Chemical Biology 3, 373–382 (2008).

31. Kramer, G. et al. L23 protein functions as a chaperone docking site on the ribosome. Nature 419, 171–174 (2002).

32. Liu, Y. L. et al. Three-Dimensional Two-Color Dual-Particle Tracking Microscope for Monitoring DNA Conformational Changes and Nanoparticle Landings on Live Cells. ACS Nano 14, 7927–7939. ISSN: 1936086X (2020).

33. Eggeling, C., Fries, J. R., Brand, L., Günther, R. & Seidel, C. A. Monitoring conformational dynamics of a single molecule by selective fluorescence spectroscopy. Proceedings of the National Academy of Sciences of the United States of America 95, 1556–1561. ISSN: 00278424 (1998).

34. König, I. et al. Single-molecule spectroscopy of protein conformational dynamics in live eukaryotic cells. Nature Methods 12, 773 (2015).

35. Di Leonardo, R., Ianni, F. & Ruocco, G. Computer generation of optimal holograms for optical trap arrays. Optics Express 15, 1913. ISSN: 1094-4087 (2007).

36. Guo, C. et al. A fast-converging iterative method based on weighted feedback for multi-distance phase retrieval. Scientific Reports 8, 1–10. ISSN: 20452322. http://dx.doi.org/10.1038/s41598-018-24666-8 (2018).

37. Guo, C. et al. Lensfree on-chip microscopy based on dual-plane phase retrieval. Optics Express 27, 35216. ISSN: 1094-4087 (2019).

38. Vestergaard, C. L., Blainey, P. C. & Flyvbjerg, H. Optimal estimation of diffusion coefficients from single-particle trajectories. Physical Review E - Statistical, Nonlinear, and Soft Matter Physics89. ISSN: 15502376 (2014).

39. Allgower, E. L. Exact inverses of certain band matrices. Numerische Mathematik 21, 279–284. ISSN: 09453245 (1973).

40. Schreiber, J. pomegranate: Fast and flexible probabilistic modeling in python. Journal of Machine Learning Research 18, 1–6. ISSN: 15337928. 1711.00137 (2018).

41. Datsenko, K. A. & Wanner, B. L. One-step inactivation of chromosomal genes in Escherichia coli K-12 using PCR products. Proceedings of the National Academy of Sciences of the United States of America 97, 6640–6645. ISSN: 00278424 (2000).

## References

9. Masullo, L. A. et al. Pulsed interleaved MINFLUX. Nano Letters 21, 840–846 (2021).

